# Importin 13-dependent Axon Diameter Growth Regulates Conduction Speeds along Myelinated CNS Axons

**DOI:** 10.1101/2023.05.19.541431

**Authors:** Jenea M Bin, Daumante Suminaite, Silvia K. Benito-Kwiecinski, Linde Kegel, Maria Rubio-Brotons, Jason J Early, Daniel Soong, Matthew R Livesey, Richard J Poole, David A Lyons

## Abstract

Central nervous system neurons have axons that vary over 100-fold in diameter. This diversity contributes to the regulation of myelination and conduction velocity along axons, which is critical for proper nervous system function. Despite this importance, technical challenges have limited our understanding of mechanisms controlling axon diameter growth, preventing systematic dissection of how manipulating diameter affects conduction along individual axons. Here, we establish zebrafish as system to investigate the regulation of axon diameter and axonal conduction *in vivo*. We identify a role for importin 13b specifically in axon diameter growth, that is independent of axonal length and overall cell size. Impairment of diameter growth along individual myelinated axons reduced conduction velocity in proportion to the diameter, but had no effect on the precision of action potential propagation or ability to fire at high frequencies. This work highlights an axon diameter-specific mechanism of growth that regulates conduction speeds along myelinated axons.

## Introduction

Across the vertebrate central nervous system (CNS), neurons have axons with incredibly diverse diameters, ranging from ∼0.1 µm to >10 µm, which corresponds to over ten thousand-fold differences in cross-sectional area ^1–3^. Axon diameter can also vary across the length of single axons ^4^, change during key developmental stages such as adolescence ^5, 6^, and may also be dynamically regulated throughout life in response to neuronal activity ^7–9^. Diversity of axon diameter is key in shaping neuronal circuit function, as it is a major determinant of the speed of action potential propagation along the axon. It has also been proposed that axons of distinct diameters may have the capacity to sustain different frequencies of action potential firing ^3^. Furthermore, biophysical and molecular signals associated with axon diameter are critical for regulating myelination along axons ^10–15^, with the presence, dimensions, and spacing of myelin sheaths all impacting conduction properties ^16^. Thus, the regulation of axon diameter contributes both directly and indirectly (via myelin) to varying and achieving the precise and rapid timings of signal transmission required for proper nervous system function. Notably, changes to axon diameter have been observed across a wide variety of neurodevelopmental, neuropsychological, and neurodegenerative diseases, and may contribute to altered network behaviour and clinical phenotypes in these patients ^17–24^. This further highlights the importance of axon diameter to the health and function of our nervous system.

Despite its importance, little is known about how axon diameter is regulated compared to other aspects of neuronal biology. The majority of studies have focussed on cytoskeletal components such as neurofilaments, microtubules, and actin-spectrin rings that influence axon diameter ^25–31^, but the cell intrinsic programmes or cell-cell interactions that lead to CNS axons having such very different sizes and functions remain largely unexplored. This is in part due to technical challenges in visualizing, measuring, and following axon diameter over time in living systems. Furthermore, along myelinated axons where multiple parameters (e.g. diameter, myelin sheath length and thickness, nodes of Ranvier, ion channels) can all impact conduction speed and covary with one another ^16, 32^, it is unclear how axon diameter growth affects conduction properties of individual axons as they develop within circuits. To address fundamental questions about axon diameter, there is a need for systematic approaches that tie together molecular manipulations with structural and functional analyses of single axons over time in living systems.

Here, we combined the strengths of zebrafish for genetic screens, high-resolution live-imaging, and electrophysiology to study the growth of axons in diameter and how this growth influences conduction along the axon. We identified a mutation in the gene encoding importin 13b (ortholog of the gene encoding human importin 13), a protein highly enriched in the nervous system ^33^, which results in a striking reduction in axon diameters. Using transgenic reporters to label specific neurons and follow their axons over time *in vivo*, we show that importin 13b function in neurons is required for the growth of axons in diameter, but not their growth in length or cell body size. The reduced diameter growth was associated with both fewer axonal neurofilaments and decreased neurofilament spacing. By combining our structural live-imaging with electrophysiological recordings from the exact same neuron, we find that the disruptions to diameter growth result in slowing of conduction velocity directly proportional to diameter, without affecting the ability to fire at high frequencies or precision of action potential firing. This implicates diameter growth as the major mechanism by which myelinated axons normally increase their conduction speed during development.

## Results

### Zebrafish as a model to study axon diameter

To investigate how CNS axons grow in diameter, we used zebrafish as a model system. Within the first five days of development in the zebrafish CNS, there is rapid and differential growth of axons in diameter, such that by 5 days post-fertilization (dpf) there is already over ∼40-fold difference in axon diameter across different neurons (Figure 1A). In particular, we focussed on the Mauthner neuron, because of its ease of identification and the fact that it has an axon that grows to a very large diameter during early nervous system development. Each fish has two Mauthner neurons, which exist as a bilateral pair within rhombomere 4 of the hindbrain and project their axons down the entire length of the contralateral spinal cord (Figure 1B). From 2 dpf (before myelination) through to 5 dpf, the Mauthner axon grows rapidly from ∼1.0 µm to ∼3.5 µm, which can readily be measured using super-resolution live-imaging (Figure 1C,D). The Mauthner axon is the first to become myelinated and becomes ensheathed along its entire length by 3.5 dpf, after which both the axon and myelin continue to grow in thickness (Figure 1D). The rapid and extensive growth in diameter of the Mauthner axon, its early myelination, and its individually identifiable nature make it a powerful model to study mechanisms of axon diameter growth, and how manipulating axon diameter affects the conduction along myelinated axons.

**Figure 1:**
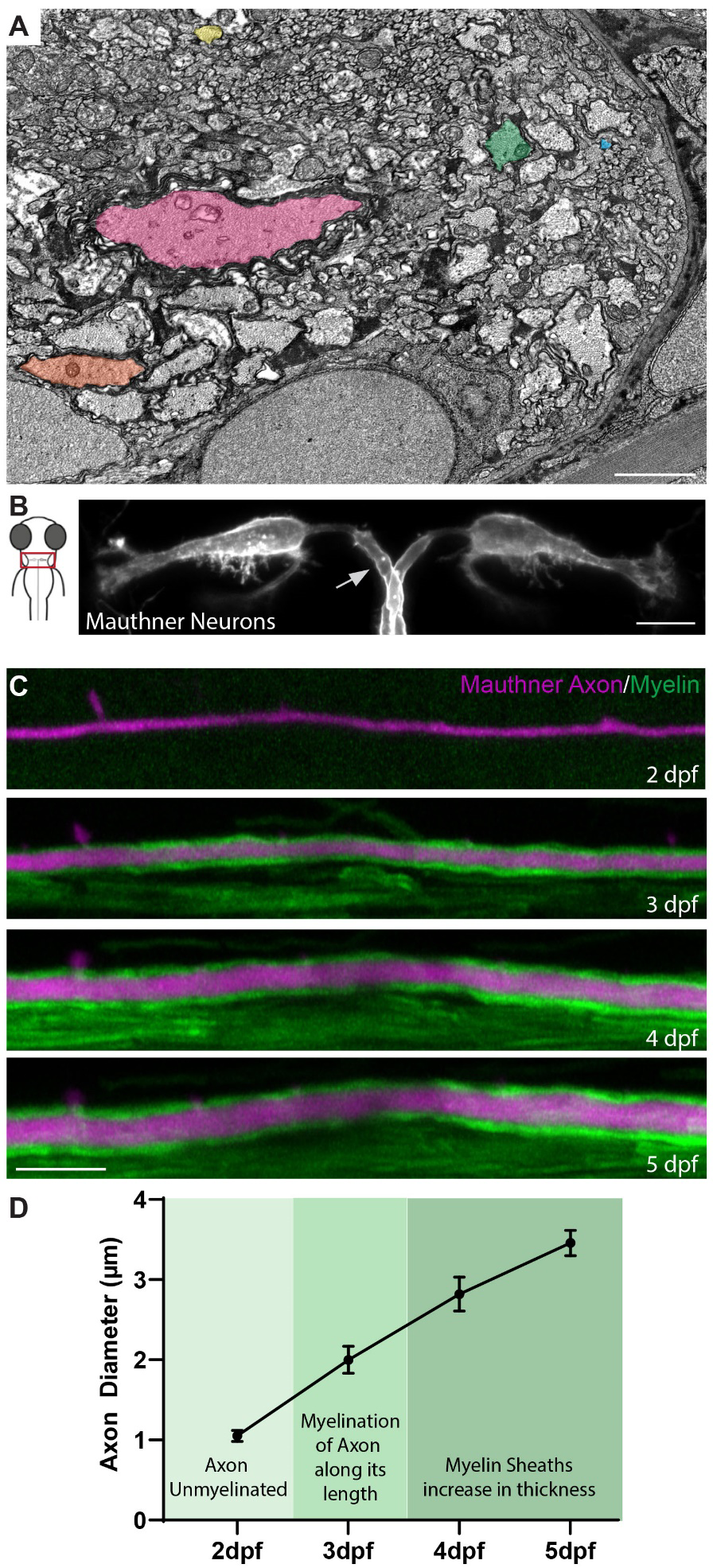
Zebrafish as a model to study axon diameter. (A) Electron micrograph of a cross section of the zebrafish ventral spinal cord at 5 dpf showing the diverse range of axon diameter. Five axons spanning the range of diameters are highlighted: the largest myelinated Mauthner axon (pink), two other myelinated axons (orange and green) and two unmyelinated axons (yellow and blue). (B) Schematic dorsal view of the larval zebrafish head with inset indicating position of Mauthner neurons show in right, labelled using the transgenic line Tg(hspGFF62A:Gal4); Tg(UAS:mem-Scarlet). Arrow points to the one of the two axons, about to cross the midline. (C) Time course depicting the growth in diameter of the Mauthner axons (magenta - Tg(hspGFF62A:Gal4); Tg(UAS:mRFP)) with myelination (green – Tg(mbp:eGFPCAAX)) (somite 15 region of the spinal cord) from 2 dpf – 5 dpf. (D) Quantification of Mauthner axon diameter growth with relation to its myelination, followed for the same axons over time at somite 15 from 2 – 5 dpf. Scale bars: 1 µm (A), 10 µm (B,C).

### Identification of the requirement of importin 13b function for axon diameter growth

Given how little is known about the regulation of axon diameter, unbiased, discovery-based screens are a powerful approach to uncover novel molecular mechanisms. As part of a zebrafish ENU (N-ethyl-N-nitrosourea) mutagenesis-based forward genetic screen aimed at identifying novel regulators of different aspects of myelinated axon development ^34–36^, we identified a mutant allele, *ue57*, which resulted in a striking reduction in the diameter of the Mauthner axon in the zebrafish spinal cord (Figure 2 A-C). The *ue57* mutant animals exhibited normal overall growth and gross morphology (Figure 2A) throughout embryonic development and early larval stages; however, they displayed reduced motility (Figure 2D) and lethality between 9-11 dpf.

**Figure 2:**
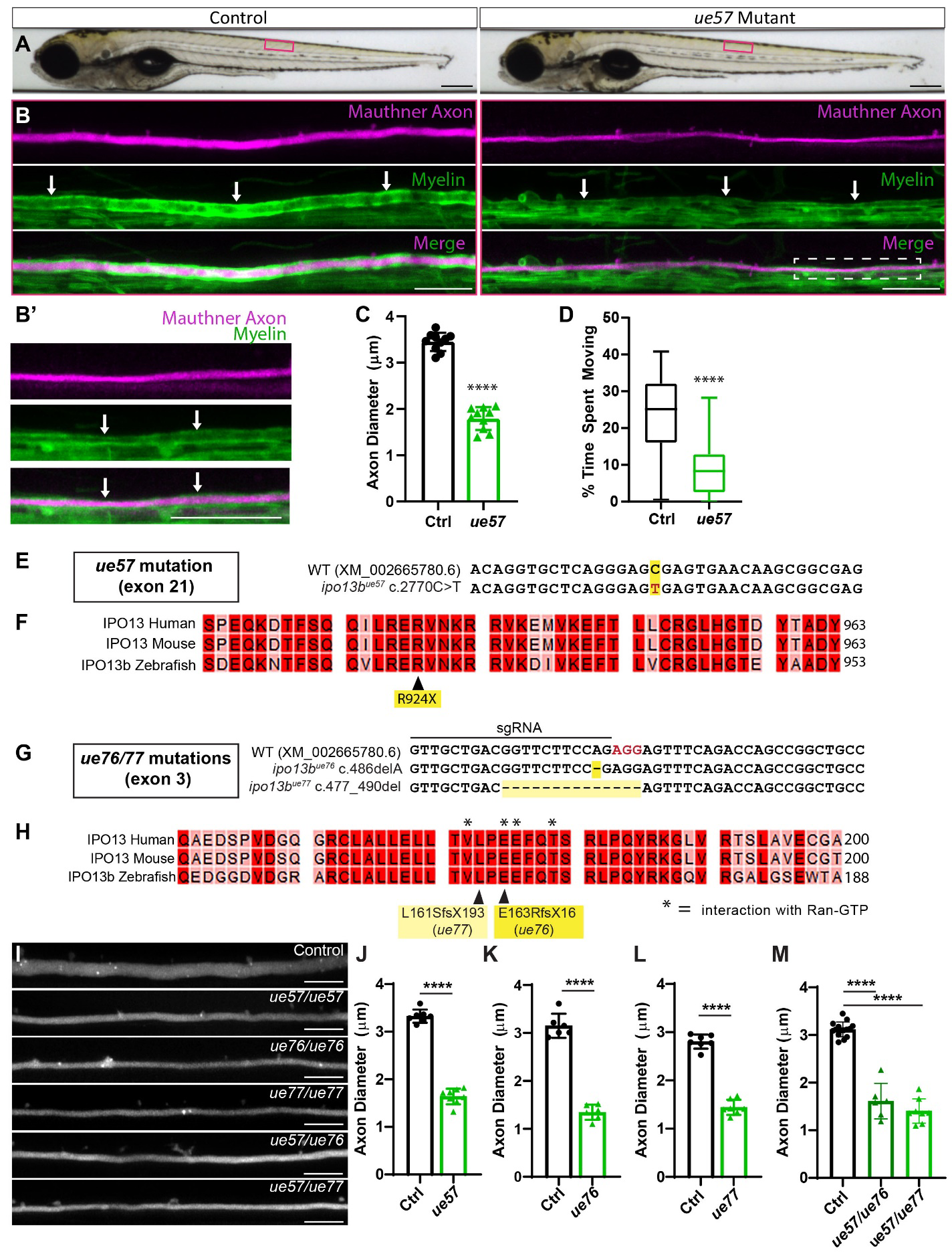
Identification of importin 13b mutants with reduced axon diameter growth. (A) Brightfield images of control and *ue57* mutant zebrafish at 5 dpf, depicting normal growth and gross morphology. (B) The Mauthner axon (somite 15) labelled using Tg(hspGFF62A:Gal4); Tg(UAS:mRFP) in control and *ue57* mutant zebrafish illustrates the smaller axon diameter in mutants at 5dpf. Labelling of myelin with Tg(mbp:eGFPCAAX) shows that the Mauthner axon is myelinated (white arrows). Magnification of the mutant Mauthner axon (region within dashed square) is shown in (B’). (C) Mauthner axon diameter measured at somite 15 using the Tg(hspGFF62A:Gal4); Tg(UAS:mRFP) reporter (Two-tailed unpaired t-test, ****p<0.0001). (D) In a 20 min open field test at 5 dpf, *ue57* mutant zebrafish spend significantly less time swimming than controls (Two-tailed Mann-Whitney test, ****P<0.0001, n= 144 ctrl, n = 31 *ue57*, from 4 different clutches of fish). (E) A region of exon 21 (last exon) of the *ipo13b* gene where a C>T base pair change (highlighted in red) was identified in *ue57* mutants. (F) This base pair change results in the introduction of a premature stop codon in the highly conserved C-terminal region of importin 13b, predicted to result in a truncated protein missing the last 30 amino acids. (G) Overview of a region in exon 3 of the *ipo13b* gene indicating the site targeted with an sgRNA for cas9-mediated DNA cleavage (PAM sequence highlighted in red), and the resulting mutations in the *ue76* and *ue77* mutant lines. (H) The mutations disrupt key residues previously shown to bind Ran-GTP (asterisks) ^43^, and result in frame shifts followed by premature stop codons. (I) Representative images of a lateral view of the Mauthner axon (somite 15) labeled using the Tg(hspGFF62A:Gal4); Tg(UAS:GFP) reporter in a 4 dpf control, *ipo13b^ue^*^57^, *ipo13b^ue^*^76^ and *ipo13b^ue^*^77^ zebrafish and 5 dpf *ipo13b^ue^*^57^*^/ue^*^77^ and *ipo13b^ue^*^57^*^/ue^*^77^ zebrafish, with quantification of axon diameter shown in (J-M) (****p<0.0001 Two-tailed unpaired t-test (J-L) or one-way ANOVA with Tukey’s multiple comparisons test (M)). For all bar graphs, each point represents an individual animal. Scale bars: 300 µm (A), 20 µm (B,B’), 10 µm (I) .

Mapping-by-sequencing of phenotypically mutant larvae localised the causative mutation to chromosome 20 (45-50 MB region, data not shown), and revealed a C > T base pair change that introduced a premature stop codon (R924X) within the last exon of the gene *ipo13b* (GRCz11:ENSDARG00000060618), predicted to eliminate the last 30 amino acids of the protein importin 13b (Figure 2 E,F). Importin 13 is a member of the importinβ superfamily, which is a family of receptors responsible for shuttling proteins between the nucleus and the cytoplasm ^37, 38^, including long-range transport roles moving cargoes between the nucleus and the dendrites/axons/synapses of neurons ^39–42^. Importin 13b is a well-conserved protein, with zebrafish importin 13b and human importin 13 sharing 83% amino acid identity. We confirmed *ipo13b* was the gene responsible for the *ue57* phenotype through CRISPR/Cas9-based generation of two additional mutant alleles (*ipo13b^ue^*^76^ and *ipo13b^ue^*^77^), which introduced frame-shift mutations in a ran-GTP binding site within the N-terminal region of the protein that is critical for importin 13 function (Figure 2 G,H) ^43^. The *ipo13b^ue^*^76^ and *ipo13b^ue^*^77^ mutants, as well as *ipo13b^ue^*^57^*^/ue^*^76^ and *ipo13b^ue^*^57^*^/ue^*^77^ trans-heterozygotes, phenocopied the Mauthner axon diameter defect observed in the *ue57* mutant (Figure 2 I-M). Thus, we concluded that *ipo13b* is the gene responsible for the axon diameter phenotypes in the *ue57* mutants, and further refer to this mutant line as *ipo13b^ue^*^57^.

### Importin 13b mutants exhibit reduced CNS axon diameter growth

To determine whether importin 13b is required for axon diameter growth and/or maintenance, we performed super resolution live-imaging of a fluorescent reporter expressed in the Mauthner neuron to measure its axon diameter over time from 2 dpf – 7 dpf. The time-course analysis revealed that while Mauthner axon diameter is initially normal in homozygous mutants at 2 dpf, growth in diameter significantly slows after this time point compared to both control and heterozygous Mauthner axons (Figure 3A,B). Notably though, Mauthner axons in mutants still grow to a diameter large enough that they become myelinated (Figure 2B,B’ and 4B). Despite the major defect in axon diameter growth, we observed no abnormalities in Mauthner axon outgrowth, with axons reaching the same length in both control and *ipo13b^ue^*^57^ animals (Figure 3C,D). We also did not detect any major effect on the size of the Mauthner cell body (Figure 3E,F). Together, these results indicate that importin 13b influences axon diameter by a mechanism that is different from those involved in regulating axon length or overall cell size.

**Figure 3:**
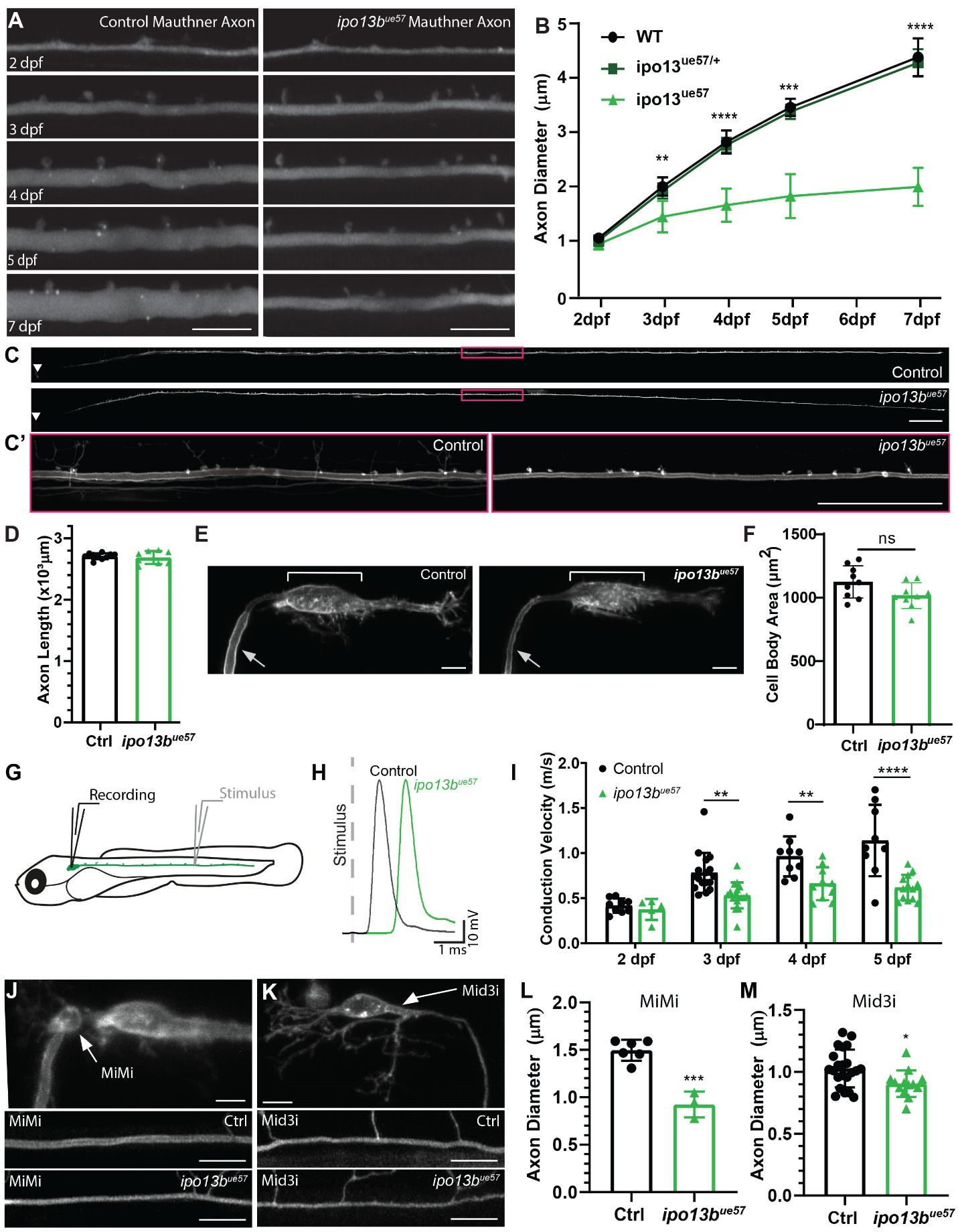
Live-imaging of axon diameter growth defects in importin 13b mutants. (A) Representative live-imaging time course of the Mauthner axon (somite 15) from 2 dpf – 7 dpf in a control and *ipo13b^ue^*^57^ zebrafish labeled using Tg(hspGFF62A:Gal4); Tg(UAS:GFP). (B) Quantification of Mauthner axon diameter growth followed for the same axons at somite 15 from 2 – 7 dpf (n=7 wildtype, 17 heterozygous, 6 mutants, 2-way ANOVA with Tukey’s multiple comparisons test, **p<0.01, ***p<0.001, ****p<0.0001, wt and het are not significantly different from one another). (C) Representative live-images of the entire Mauthner neuron in control and *ipo13b^ue57^*zebrafish at 3 dpf, which were used to measure axon length in D. Area boxed in magenta is enlarged in (C’). (D) Quantification of the entire length of the Mauthner axon at 3 dpf (Two-tailed unpaired t-test with Welch’s correction, p=0.5992). (E) Representative live-imaging of the Mauthner neuron at 4 dpf in control and *ipo13b^ue^*^57^ zebrafish labelled using the transgenic line Tg(hspGFF62A:Gal4); Tg(UAS:mem-Scarlet). Bracket indicates the position of the cell body and arrow points to the axon. (F) Quantification of the Mauthner cell body area at 4 dpf (Two-tailed unpaired t-test, p=0.0638). (G) Schematic overview of the electrophysiological set-up for measuring conduction velocity along the Mauthner axon. (H) Example traces of Mauthner whole-cell current clamp recording showing action potentials generated in response to Mauthner axon extracellular stimulation. There is a longer latency between the stimulus artifact and action potential peak in the mutants indicating a slower conduction velocity. (I) Conduction velocity measurements along Mauthner axon from 2 dpf to 5 dpf in *ipo13b^ue^*^57^ animals and control siblings (2-way ANOVA with Sidak’s multiple comparisons test, **p=0.0025 (3dpf), 0.0071 (4dpf), ****p<0.0001). MiMi (J) or Mid3i (K) neuron (top panel) and its axon (somite 15) in control (middle panel) and *ipo13b^ue^*^57^ (bottom panel) animals labelled using the transgenic reporter Tg(hspGFF62A:Gal4); Tg(UAS:mem-Scarlet). (L) Quantification of MiMi axon diameter at 5 dpf (Two-tailed unpaired t-test, ***p<0.0003). (M) Quantification of Mid3i axon diameter at 5 dpf (Two-tailed unpaired t-test, *p<0.016). For all bar graphs, each point represents an individual animal. Scale bars: 10 µm (A, E, J, K), 100 µm (C), 50 µm (C’).

We next measured conduction velocity along the *ipo13b^ue57^*mutant Mauthner axons. Action potentials were recorded by making whole-cell current clamp recordings from the Mauthner neuron cell body while delivering stimuli to its axon using an extracellular electrode (Figure 3G,H). We observed that conduction velocity along Mauthner axons in *ipo13b^ue^*^57^ mutants was slower than in controls from 3 dpf onwards, in line with the timeline of impaired axon diameter growth (Figure 3H,I). By 5 dpf, action potentials were being propagated at almost half the speed of control Mauthner axons.

To assess whether the deficits in axon diameter growth occurred in other types of neurons, we first transgenically labelled and live-imaged individual axons from two additional and easily identifiable pairs of reticulospinal neurons that project axons along the ventral spinal cord, the MiMi and Mid3i neurons. Both sets of neurons also exhibited significantly reduced axon diameter in *ipo13b^ue^*^57^ mutants at 5 dpf (Figure 3J-M). The severity of the phenotype correlated with axon diameter, such that growth of the larger diameter axons was more affected than the smaller diameter axons (growth in diameter reduced by 47% for Mauthner axons, 39% for MiMi axons, and 13% for Mid3i axons at 5 dpf).

Next, we assessed axon diameter in cross sections of the 7 dpf zebrafish spinal cord using electron microscopy (Figure 4A,B). We measured a 51% reduction in Mauthner axon diameter (80% reduction by area), which was comparable to the 55% reduction observed at 7 dpf using live-imaging (Figure 4C vs 3B). The average axon diameter of the next 30 largest axons per ventral spinal cord hemi-segment was also significantly reduced, with binning of these axons based on their axon diameter showing a global shift to smaller diameters in *ipo13b^ue^*^57^ mutants (Figure 4D,H). There was also a small, but significant, shift towards smaller axon diameters amongst the 30 largest axons in the dorsal spinal cord (Figure 4 E,F,I). As dorsal axons are on average smaller in diameter than ventral axons, the smaller effect on dorsal axons is in agreement with our live-imaging results indicating that larger diameter axons are more significantly affected by loss of importin 13b function. Our electron microscopy analyses also revealed that fewer axons were ensheathed with myelin in the spinal cord of *ipo13b^ue^*^57^ mutants compared to their sibling controls (Figure 4G). Given the role of axon diameter in the selection of axons that become myelinated ^10–14^, the reduction in myelin is likely at least in part a consequence of the reduction in axon diameter. Together, our live-imaging and electron microscopy studies support the conclusion that importin 13b is required for the proper growth of axon diameter for a wide range of neurons.

**Figure 4:**
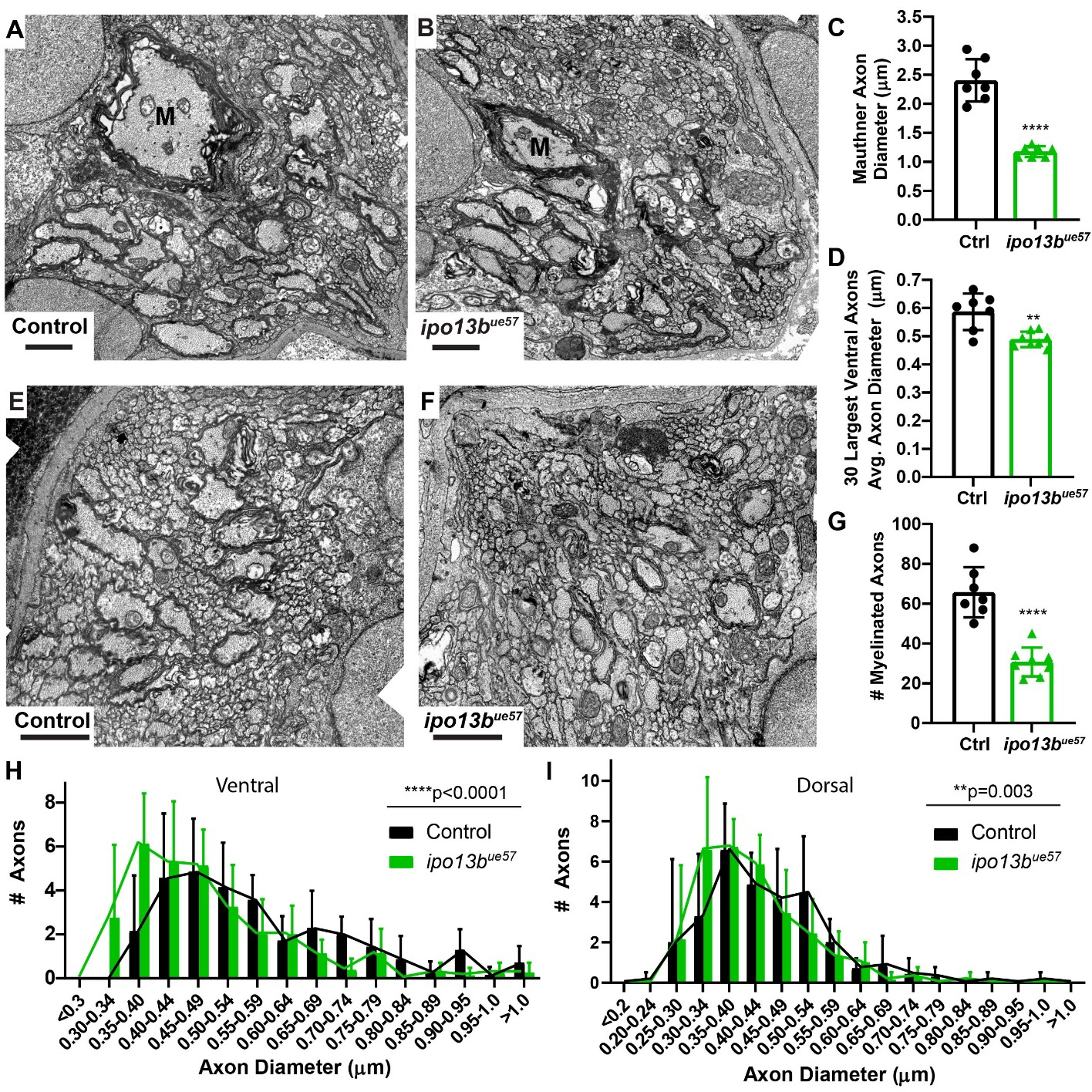
Electron microscopy of axon diameter growth defects in importin 13b mutants. (A) Representative electron micrographs of cross sections of the ventral spinal cord at 7 dpf in control and (B) *ipo13b^ue^*^57^ animals. The Mauthner axon is labelled ‘M’. (C) Quantification of Mauthner axon diameter from 7 dpf electron micrographs (Two-tailed unpaired t-test with Welch’s correction, ****p<0.0001). (D) Mean diameter for the 30 largest axons in each hemi ventral spinal cord at 7 dpf, excluding Mauthner (Two-tailed unpaired t-test with Welch’s correction, **p=0.0061). (E) Representative electron micrographs of cross sections of the dorsal spinal cord at 7 dpf in control and (F) *ipo13b^ue^*^57^ zebrafish. (G) Number of myelinated axons in the dorsal and ventral tracts of each hemi spinal cord at 7 dpf (Two-tailed unpaired t-test, ****p<0.0001). (H) Distribution of axon diameters for the 30 largest axons in each hemi ventral spinal cord at 7 dpf, excluding Mauthner (n=7 control and 8 mutants, the distributions are significantly different, Kolmogorov-Smirnov test, ****p<0.0001). (I) Distribution of axon diameters for the 30 largest axons in each hemi dorsal spinal cord at 7 dpf (n=7 control and 8 mutants, distributions are significantly different, Kolmogorov-Smirnov test, **p=0.0030). Scale bars = 1 µm.

### Loss of Importin 13 affects both neurofilament number and spacing

To better understand how loss of importin 13b function affects axon diameter growth, we looked at the axonal cytoskeleton within mutant Mauthner axons. Previous studies have implicated regulation of neurofilament (the neuron-specific intermediate filaments) expression and spacing as major contributors to the radial growth of large diameter axons (reviewed in ^44^). We analyzed the density of neurofilaments in cross sections of 7 dpf control and *ipo13b^ue^*^57^ mutant Mauthner axons in electron microscopy images (Figure 5A-B) and found that there a 34% increase in neurofilament density in *ipo13b^ue^*^57^ mutant Mauthner axons (Figure 5C). Likewise, nearest neighbour analysis revealed a shift towards shorter distances between individual neurofilaments (Figure 5D). However, given the roughly 80% reduction in cross-sectional area of *ipo13b^ue^*^57^ mutant Mauthner axons, one would have expected a 4-5x increase in neurofilament density should the number of neurofilaments have remained the same. Thus, both the number and spacing of neurofilaments is reduced when importin 13b function is disrupted.

**Figure 5:**
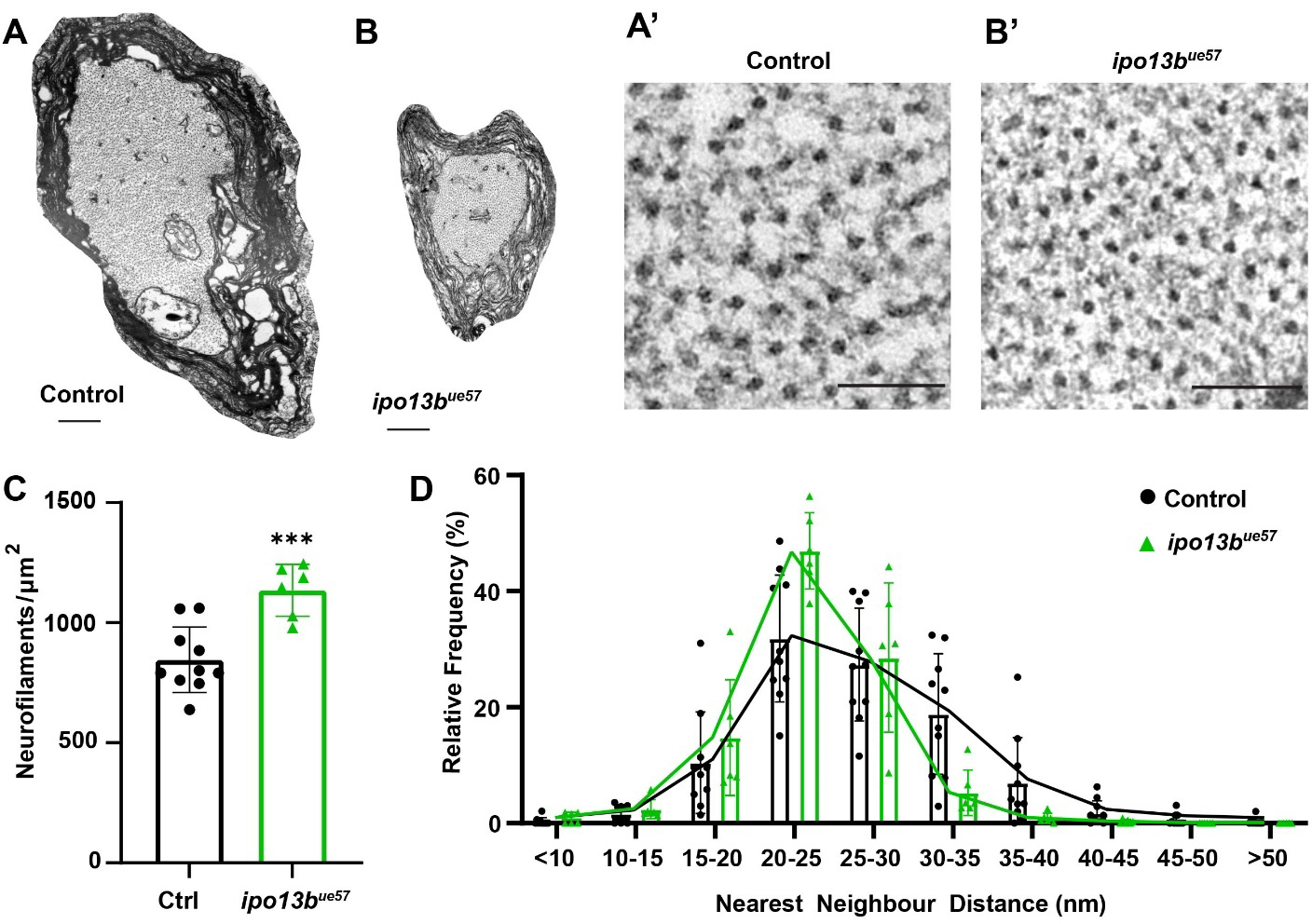
Disruption of importin 13b function alters neurofilaments in the Mauthner axon. (A-B) Representative electron micrographs of a cross section of the Mauthner axon in the ventral spinal cord at 7 dpf in control and *ipo13b^ue57^*zebrafish. A region of each axon is magnified in A’ and B’ to allow visualization of the distribution of neurofilaments. (C) Density of neurofilaments in control and *ipo13b^ue^*^57^ mutant Mauthner axons at 7 dpf (Two-tailed unpaired t-test, ***p=0.0006). (D) Nearest neighbour distribution of neurofilaments in the Mauthner axon at 7 dpf, showing a shift to closer nearest neighbours in the *ipo13b^ue^*^57^ mutants. For all graphs, each point represents an individual animal. Scale bars: 500nm (A), 100nm (B).

### Axon diameter growth requires importin 13b function in neurons

*Ipo13* is known to be expressed in all CNS cell types^45^. Thus, importin 13b may be required within neurons for their axons to grow in diameter, or it may play a role in other non-neuronal cells and exert its effect on axon diameter in a non-cell autonomous manner ^15, 46–50^. To investigate the role of importin 13b specifically in neurons, we made neuron-specific *ipo13b* mutants using a cell-type specific CRISPR/Cas9 strategy (Figure 6A, Materials and Methods). Similar to the *ipo13b^ue^*^57^ mutants, neuron-specific *ipo13b* mutants exhibited significantly reduced Mauthner axon diameter growth (Figure 6B,C). A small proportion of fish (∼20%) expressing both neuronal cas9 and sgRNAs targeting *ipo13b* showed no observable defects in the diameter of one or both Mauthner axons. Given the mosaic nature of the mutations introduced using the CRISPR/Cas9 strategy (see Materials and Methods), we expect this likely reflects the introduction of in-frame or silent *ipo13b* mutations within these neurons that do not cause bi-allelic loss-of-function. In affected Mauthner axons, the reduction in axon diameter was not as severe as in full *ipo13b* mutants (Figure 2C vs Figure 6C), which may reflect the timing or extent of loss of importin 13b function, or additional non-cell autonomous contributions of importin 13b. Together, our results indicate that importin 13 function in neurons is important for axons to grow correctly in diameter.

**Figure 6:**
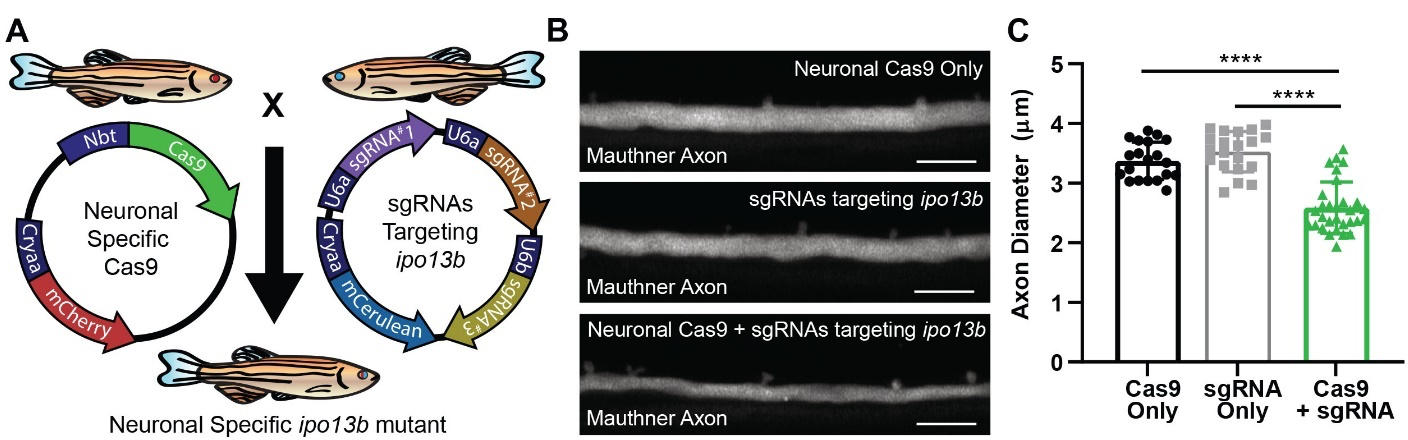
Disruption of axon diameter in neuronal specific *ipo13b* mutants. (A) Schematic overview of the transgenic CRISPR/Cas9 strategy used to generate neuron specific *ipo13b* mutants. (B) Representative images of the Mauthner axon (somite 15) in the neuron specific *ipo13b* mutants (cas9+sgRNA) and control animals (cas9 or sgRNA only) labelled using Tg(hspGFF62A:Gal4); Tg(UAS:mRFP). (C) Quantification of Mauthner axon diameter in the neuron specific *ipo13b* mutants (Kruskal-Wallis test with Dunn’s multiple comparisons test, ****p<0.0001). Each point represents an individual Mauthner axon. Scale bars = 10 µm.

### The growth of axons in diameter drives dynamic changes to conduction along myelinated axons

We next asked how axon diameter growth affects conduction along a single myelinated axon in an intact neural circuit *in vivo*. Previous studies comparing myelinated axons of different diameters have found that there is a linear relationship between axon diameter and conduction velocity, whereby increasing axon diameter results in increased action potential speed ^16, 51^. However, it is not clear how conduction velocity is controlled along individual myelinated axons over time *in vivo*. Many other features of the myelinated axon could be regulated independently of axon diameter to alter conduction, such as the length and thickness of myelin sheaths, nodal dimensions, and the number, type, and density of ion channels ^16, 32^. Thus, we wanted to ask: “what happens to conduction when axon diameter growth is disrupted?” Would the linear relationship between axon diameter and conduction velocity be maintained or would it be altered by regulation of other features of the myelinated axon? If axon diameter was the driver of the changes to conduction velocity over time, we would expect control and *ipo13b* mutant axons of the same diameter to propagate action potentials at the same speed, despite being different developmental ages. If other features of the myelinated axon were regulated independently of axon diameter, the speed of action potential propagation in *ipo13b* mutants might significantly deviate from the expected linear relationship between axon diameter and conduction velocity.

To address the relationship between axon diameter growth and action potential speed along individual axons, we performed live-imaging to determine the diameter of individual Mauthner axons, and then carried out electrophysiological recordings from the same neuron. We first assessed how conduction velocity relates to axon diameter along myelinated control axons as they grew between 3 dpf and 5 dpf. As has previously been observed when comparing myelinated axons of different diameters ^16, 51^, conduction velocity increased in a linear relationship with axon diameter (Figure 7A-C).

**Figure 7:**
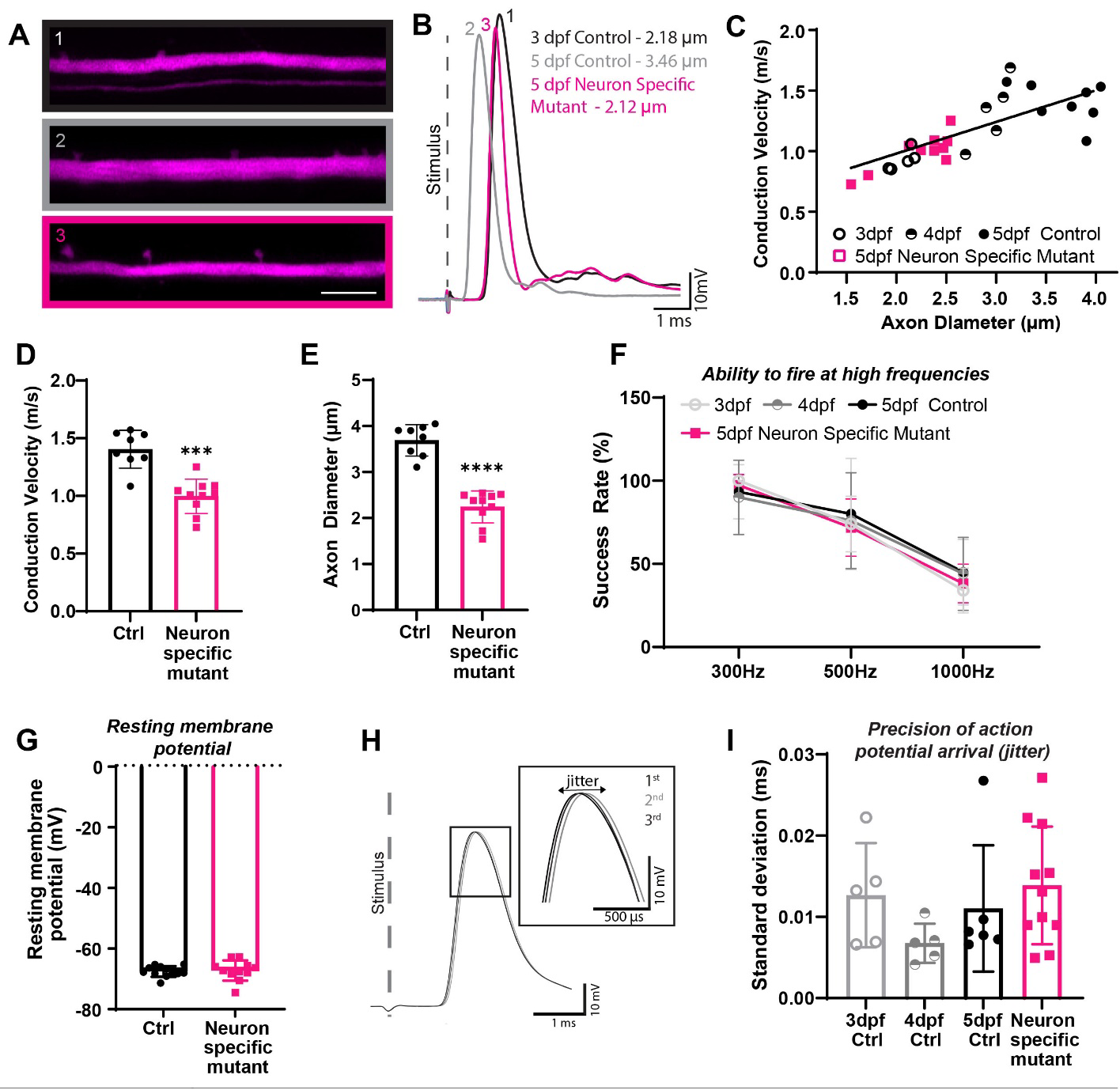
Diameter growth drives changes to conduction speeds along myelinated axons over time. (A) Super-resolution imaging of Mauthner axons labelled using the transgenic line Tg(hspGFF62A:Gal4); Tg(UAS:RFP) alongside whole-cell patch clamp traces showing action potentials generated in response to extracellular stimulation of the same axons (B). (C) Conduction velocity along Mauthner axon plotted against axon diameter for controls axons (3-5dpf) and neuronal specific *ipo13b* mutant axons (5 dpf). Control points are fitted with a linear regression line (R=0.4959 for control, neuronal specific ipo13 mutants are not significantly different from controls by simple linear regression test). (D) Conduction velocity measurements along Mauthner axon at 5 dpf in neuronal specific *ipo13b* mutant animals and control siblings (Mann-Whitney Test, ***p=0.0002). (E) Axon diameter measurements for the same axons as in (D) (Mann-Whitney Test, ****p<0.0001). (F) The success rate of action potential firing by the Mauthner neuron in response to 10 stimulations at 300 Hz, 500 Hz and 1000 Hz (no significant differences using mixed-effects analysis with Sidak’s multiple comparisons test, n=5 3dpf controls, 5 4dpf controls, 6 5dpf controls, 10 *ipo13b^ue^*^57^ mutants, 11 neuronal specific *ipo13b* mutants). (G) Resting membrane potential in control and neuronal specific *ipo13b* mutant Mauthner neurons (no significant differences, Two-tailed unpaired t-test). (H) Depiction of three consecutive action potentials, which have slight variations in their latency of arrival. This variation is referred to as jitter. (I) The precision of action potential arrival (jitter) along Mauthner axon (no significant differences, one-way ANOVA with Tukey’s multiple comparisons test). For all bar graphs, and C, each point represents an individual animal. Scale bars = 10 µm.

Next, we asked whether conduction velocity along importin 13b mutant Mauthner axons followed this same relationship between axon diameter and action potential speed, or whether they deviated from what was observed in controls, indicative of diameter-independent changes that affect conduction. For this experiment, we used neuron-specific *ipo13b* mutants, rather than the full *ipo13b^ue^*^57^ mutants, to exclude possible effects on conduction due to importin 13b autonomous function in oligodendrocytes on myelin formation. Strikingly, we found that while the neuron-specific *ipo13b* mutant Mauthner axons conducted action potentials slower than control axons of the same development stage, conduction speeds along mutant axons were comparable to control axons of the same axon diameter at earlier developmental stages (Figure 7A-E). This indicates that increases in conduction velocity along the myelinated Mauthner axon are primarily driven by its growth in diameter. It also suggests that during this developmental window, the axon does not adapt (e.g. by modulating myelin or ion channels) to compensate for its reduced diameter in order to achieve its normal conduction speed.

### Reduced axon diameter does not affect the ability to fire at high frequencies

In addition to its influence on conduction velocity, it has also been speculated that larger diameter axons better support higher frequency firing of action potentials ^3^. Therefore, we asked whether the ability of the Mauthner axon to sustain high-frequency firing was influenced by its diameter. To test this, we examined the action potential activity evoked by a range of depolarising stimuli delivered at 300 Hz, 500 Hz and 1000 Hz to Mauthner axons in control and *ipo13b^ue57^*mutant animals. Despite being much smaller in diameter than controls, *ipo13b^ue^*^57^ mutant axons maintained even the highest frequency trains of stimuli (Figure 7F). This counters the view that axon diameter is a parameter that is specifically regulated to sustain high-frequency firing patterns, at least along myelinated axons. Interestingly, we recently reported that hypomyelination of the Mauthner axon does affect its ability to fire at high frequencies ^52^. Thus, these results indicate that it is the myelin on large diameter axons that is important to support high frequency firing, rather than their diameter itself. In addition, we also did not detect any differences in the resting membrane potential of the *ipo13b^ue57^*mutant Mauthner neurons indicating normal intrinsic excitability (Figure 7G), nor in the precision of action potential propagation (jitter) (Figure 7H,I). Together, our data indicates disrupting axon diameter growth specifically affects the speed of action potential propagation, without affecting intrinsic excitability, the ability to fire at high frequencies or precision of action potential arrival.

## Discussion

Here we exploited the tractability of the zebrafish for genetics, *in vivo* imaging, and electrophysiology, to study how axons grow in diameter and how axon diameter growth affects the dynamic changes in conduction properties observed along individual myelinated axons over time *in vivo*. We demonstrate that the nuclear transport receptor importin 13b is essential for axon diameter growth via a mechanism that affects both neurofilament number and spacing, but that is regulated independently of axon outgrowth or growth the neuron as a whole. To examine the relationship between axon diameter growth and conduction velocity during development, we combined super-resolution live-imaging with electrophysiology to assess axon diameter and conduction velocity along individual myelinated axons. In doing so, we show that axon diameter growth defines the conduction speed of myelinated axons, but is not required to sustain high frequency firing or for precision of action potential propagation. Together, this highlights the involvement of importin 13b in an axon diameter-specific neuronal growth mechanism that regulates conduction speeds along myelinated axons.

A huge advantage of our live-imaging paradigm is the is the ability to image single neurons in their entirety, allowing us to assess not only changes to axon diameter over time, but also how this relates to other aspects of neuronal growth. This allowed us to determine that while importin 13b neurons have axons with smaller diameter axons, their axon length and cell body size are unaffected. There is limited knowledge of the molecular pathways that regulate axon diameter growth and as such, previous manipulations of axon diameter have relied on targeting global growth pathways within the neuron (e.g. AKT pathway) that also affect axon diameter ^11, 53–55^. Thus, importin 13b offers a molecular entry point to studying the specific control of axon diameter and how this influences conduction along myelinated axons.

The reduction in axon diameter in importin 13b mutants was associated with both reduced neurofilament number and spacing within the axon. Neurofilaments are the major cytoskeletal component controlling axon diameter, but the molecular mechanisms controlling their expression and post-translational modifications that influence diameter remain poorly understood. Therefore, extensive future studies will be required to uncover the molecular pathway(s) in which importin 13b functions to impact neurofilaments and axon diameter. Over 600 candidate cargo for importin 13 have been identified, ranging from transcription factors to proteins that directly contribute to or modify the axonal cytoskeleton^56–59^. In Drosophila, importin 13 has been shown to localize at the synapse ^60^ raising the additional possibility that importin 13b could be involved in the transport of activity-related or target-derived signals, both of which have previously been suggested to modulate of axon diameter ^7–9, 15^.

While axon diameter is a key parameter controlling the speed of action potential propagation along myelinated axons, conduction velocity is also influenced by several other factors including the length and thickness of myelin sheaths, the length of the nodes of Ranvier between myelin sheaths, and the composition, clustering, and density of ion channels along the axon ^16, 32^. Many of these parameters often covary with one another as myelinated axons grow in diameter over time *in vivo*. For example, as myelinated axons grow in diameter their myelin sheaths will often increase in thickness, and it is thought that this is at least in part regulated by axon diameter itself ^8, 15^. However, it is unclear the extent to which developmental changes along the myelinated axon that impact conduction are driven by axon diameter growth versus independently regulated over time. We took advantage of the fact that we could compare the conduction along myelinated axons from the same neuron at the same development time point, but with significantly different axon diameter. Our results demonstrate that conduction velocity along the smaller diameter Mauthner axons was precisely scaled to their diameter, such that they had the same conduction velocity observed along younger control axons of the same diameter. Had other parameters along individual axons (e.g. myelin) been regulated independently of diameter to modulate conduction speeds, we might have expected importin 13b mutant axons to exhibit much faster conduction velocities than observed for control axons of the same diameter. Therefore, our data indicates that the changes to conduction velocity observed along myelinated axons during this developmental time window are driven by their growth in diameter. This underscores the need to better understand the regulation of axon diameter as a mechanism to control the timing of action potential propagation within neuronal circuits.

Although axon diameter dictated conduction velocity speeds along individual myelinated axons, other aspects of conduction, including the ability to sustain high frequency firing and precision of action potential arrival, were independent of axon diameter and thus likely set by other parameters of the myelinated axon. Going forward a major challenge for the field will be to move towards a holistic view of myelinated axon structure and function, which will require an understanding of the regulation of each aspect of the entire myelinated axon unit (e.g. axon diameter, myelin, nodal dimensions, ion channel composition) individually and in relation to one another. In recent years a great deal of attention has been placed on the fact that myelination by oligodendrocytes responds to neuronal activity and how this may in turn dynamically modulate neural circuit function; for recent reviews see ^32, 61–63^. However, recent studies have shown that axon diameter can also be influenced by neural activity ^7, 8^, suggesting that changes to axon diameter may underlie some of the activity-dependent changes to myelin. It could be argued that the dynamic regulation of axon diameter provides a much simpler means to set and adjust the conduction properties of myelinated axons, compared to regulating numerous independent myelin sheaths, or axonal sub-domains, along the entire length of a myelinated axon. Nonetheless, future studies will be required to determine when and to what extent each of the function-regulating aspects of myelinated axons can change over time, which aspects co-vary and which can be independently regulated. The fact that we have validated the use of zebrafish as a model in which changes to axon diameter can be both identified and interrogated means that we can use zebrafish as a system to experimental disentangle how distinct components of the myelinated axon collaborate to determine the function of the entire myelinated axon unit.

## Acknowledgements

We thank members of the Lyons laboratory for feedback and the University of Edinburgh BVS Zebrafish Facility, Zebrafish Imaging and Screening Facility, and Transmission Electron Microscopy Facility for expert assistance. This work was supported by Wellcome Trust Senior Research Fellowships (102836/Z/13/Z and 214244/Z/18/Z), and a Lister Institute Research Prize to D.A.L. J.M.B was supported by a Canadian Institute of Health Research a postdoctoral fellowship and a Multiple Sclerosis Society of Canada/ Fonds de la recherche du Quebec-Sante postdoctoral fellowship.

## Material and Methods

### RESOURCE AVAILABILITY

#### Lead Contact

Further information and requests for resources and reagents should be directed to and will be fulfilled by the Lead Contact, David Lyons.

#### Materials Availability

Materials generated in this study are available by contacting the lead contact.

#### Data and Code Availability

The code generated to measure axon diameter is available at https://github.com/jasonjearly/Axon_Caliber. The published article includes all datasets generated and analyzed in this study.

### EXPERIMENTAL MODELS AND SUBJECT DETAILS

#### Zebrafish husbandry and transgenic/mutant lines

Adult zebrafish were housed and maintained in accordance with standard procedures in the Queen’s Medical Research Institute zebrafish facility, University of Edinburgh. All experiments were performed in compliance with the UK Home Office, according to its regulations under project licenses 60/4035, 70/8436, and PP5258250. Adult zebrafish were subject to a 14/10 hr, light/dark cycle. Embryos were produced by pairwise matings and were staged according to days post-fertilisation (dpf). Embryos were raised at 28.5°C, with 50 embryos (or less) per 10 cm petri dish in ∼45mL of 10 mM HEPES-buffered E3 embryo medium or conditioned aquarium water with methylene blue. All experiments used zebrafish larvae between 3 – 7 dpf, on a wildtype (AB/WIK/TL) or nacre^−/−^ ^64^ background. At these ages, sexual differentiation of zebrafish has not yet occurred.

Throughout the text and in figures, ‘Tg’ denotes a stable, germline inserted transgenic line. The following lines were used in this study: Tg(mbp:EGFP-CAAX) ^65^, Tg(hspGFF62A:Gal4) ^66, 67^, Tg(UAS:mRFP) ^66^, Tg(UAS:GFP) ^66^, Tg(UAS:mem-Scarlet) (generated in this study). The *ipo13b^ue^*^57^ mutant was identified during an ENU-based forward genetic screen, described below. The *ipo13b^ue^*^76^ and *ipo13b^ue^*^77^ mutants were generated using CRISPR/Cas9 based gene targeting, as described below.

### METHOD DETAILS

#### ENU Mutagenesis Screen

The ENU mutagenesis screen in which the *ipo13b^ue^*^57^ mutant was generated was previously published ^35, 36^. Briefly, 10 adult AB males were mutagenized with 3.5mM ENU for 1 hr per week over three consecutive weeks. Assessment of mutagenesis efficiency was made by crossing with carriers of a pigment mutation, *sox10^cls^* ^68^, and well-mutagenised males were crossed with AB females to generate the F1 generation. F1 individuals were bred with Tg(mbp:GFP-CAAX) animals to introduce a myelin reporter into the mutagenized stocks and generate individual F2 families. In total, 212 F2 families were generated, from which 946 clutches were screened for disruption to normal myelin profiles observed at 5 dpf. For extensive protocol details on zebrafish ENU-based mutagenesis forward genetic screens, see ^34^.

#### *ue57* mapping-by-sequencing

Following an outcross to WIK, pooled DNA from mutant recombinants and non-mutant recombinants was sequenced separately on a Illumina HiSeq4000 (Edinburgh Genomics). We processed this data through a modified version of the Variant Discovery Mapping CloudMap pipeline ^69^, on an in-house Galaxy server using the Zv9/danRer7 genome and annotation. For both the Variant Discovery Mapping plots and assessing the list of candidate variants we subtracted a list of wildtype variants compiled from sequencing of the EKW strain plus previously published data ^70–72^. The mapping identified a 5 Mb mapping region on Chr 20 between 45-50Mb. From the variant list within this region, we filtered for prospective missense and nonsense mutations likely to result in strong loss of function of encoded proteins. This candidate list was further filtered by excluding polymorphisms found in other mutants that we sequenced that derived from the same ENU screen. This revealed seven protein coding variants within the region, of which only the variant in *ipo13b* was a predicted nonsense mutation.

#### *ue57* genotyping

*ue57* fish were genotyped as follows. A 205 bp PCR product including the *ue57* mutation was amplified using the following primers: Fwd 5’ – CCACTAATAAAGACTATGTTTTCTCCTTTTT**G**CTC and Rev 5’ – CTTCAGTTTCCGACACAGTATTG. Note, the bolded G in the fwd primers represents of T>G mismatch introduced to generate an MwoI restriction enzyme cut site in the wildtype DNA sequence. 10 µM of each primer was combined with OneTaq QuickLoad Master Mix (NEB #M0486) and run in a thermocycler as follows: 94°C for 2 min; 36 cycles of 94°C for 30 s, 51°C for 20 s, 68°C for 10 s; and 68°C for 5 min. The PCR product was digested with 1 unit of MwoI (NEB #R0573S) for 2 hrs at 60°C, then run on a 3% agarose gel at 150 V for 45 min. The *ue57* mutation disrupts the MwoI cut site resulting in a single DNA band of ∼205 bp, while wildtype fish have two DNA bands of 174 bp and 31 bp.

#### CRISPR/Cas9 based targeting

To independently disrupt *ipo13b* function, we designed an sgRNA (GTTGCTGACGGTTCTTCCAG) targeting a region in exon 3 previously shown to interact with ran-GTP ^43^, the binding to which is essential for importin 13 function. The sgRNAs were generated using the MEGAshortscript T7 Transcription Kit (Invitrogen #AM1354) according to manufacturer’s instructions and purified using MEGAclear Transcription Clean-up Kit (Invitrogen #AM1908) according to manufacturer’s instructions. To synthesize the template DNA required for the sgRNA in vitro transcription, a two-oligo PCR method was used, as described in ^73^. The *ipo13b*-specific oligo sequence was 5’-AATTAATACGACTCACTATA<u>GGTGCTG ACGGTTCTTCCAG</u>GTTTTAGAGCTAGAAATAGC. The scaffold oligo sequence was 5’-GATCCGCACCG

ACTCGGTGCCACTTTTTCAAGTTGATAACGGACTAGCCTTATTTTAACTTGCTATTTCTAGCTCTAAAAC. The PCR reaction was performed using Phusion High-Fidelity DNA Polymerase (NEB #M0530) with 10 µM of each primer (synthesized by IDT) and run in a thermocycler as follows: 95°C for 2 min; 30 cycles of 95°C for 10 s, 60°C for 20 s, 72°C for 10 s; and 72°C for 5 min. The full-length PCR product was purified using Monarch Gel Extraction Kit (NEB #T1020), according to manufacturer’s instructions.

500 ng of sgRNA was combined with 20 µM EnGen Cas9 NLS (NEB #M0646) in 1x Cas9 Nuclease Reaction Buffer (NEB #B0386A) (total volume 3 µl) and incubated at 37°C for 5 min, then transferred to ice. Approximately 1 nL of the sgRNA-cas9 ribonucleoprotein complex was injected into Tg(mbp:EGFP-CAAX) embryos at the one-cell stage. Embryos were raised to adulthood and outcrossed to nacre^−/−^ fish to generate the F1 generation. F1 fish were screened for the presence of CRISPR/Cas9 induced mutations by amplifying a region of exon 3 using the primers 5’-GACTCCCTCAAATCCCAGCTC and 5’ – GACAGACACTTGAGCACACG. 10 µM of primers were combined with OneTaq QuickLoad Master Mix (NEB #M0486) and run in a thermocycler as follows: 94°C for 2 min; 30 cycles of 94°C for 30 s, 57°C for 30 s, 68°C for 30 s; and 68°C for 5 min. The 385 bp PCR product was digested with 1 unit of Hpy188III (NEB #R0622) for 2 hrs at 37°C, then run on a 2% agarose gel at 180V for 45 min. Mutations disrupted a Hpy188III cut site resulting in a DNA band of ∼231 bp, rather than DNA two bands of 162 bp and 69 bp. The exact indel size and location was determined by Sanger sequencing of the 385 bp PCR product following by analysis using CrispID crisped.gbiomed.kuleuven.be, ^74^ to de-convolute the overlapping spectra from the wildtype and mutant alleles. Two different mutations (*ipo13b^ue^*^76^, c.486delA and *ipo13b^ue^*^77^, c.477_490del) were selected to propagate F2 generations, which were then in-crossed to generate the mutants used in this study. Both mutations result in frameshifts and premature stop codons.

#### Generation of UAS:mem-Scarlet Transgenic Line

The UAS:mem-Scarlet expression vector was generated using Gateway cloning. mScarlet was PCR amplified from the plasmid pmScarlet_C1 (Addgene plasmid #85042) while also adding a fyn myristolyation sequence (a membrane targeting sequence) to the 5’ end using the following primers: mem-Scarlet_F GGGGACAAGTTTGTACAAAAAAGCAGGCTGCCACCATG<u>GGCTGTGTGCAATGTA</u> <u>AGGATAAAGAAGCAACAAAACTGACG</u>GTGAGCAAGGGCGAGGCAG 3’ and mem-Scarlet_R GGGGACCA CTTTGTACAAGAAAGCTGGGT-TTACTTGTACAGCTCGTCCATGCCG. A middle entry vector (pME_mem-Scarlet) was generated using 150 ng of purified PCR product with 150 ng of pDONR221in a BP reaction using BP clonase II performed overnight at room temperature. To generate the final Tol2 expression vector (UAS:mem-Scarlet), 20 fmol of each of the following entry and destination vectors were combined in a LR reaction using LR Clonase II Plus performed overnight at room temperature: p5E_10UAS (Tol2Kit #327) ^75^, pME_mem-Scarlet, p3E-polyA (Tol2Kit #302), and pDestTol2pA2-cryaa:mCherry (Addgene #64023). Note, the destination vector includes the red eye marker cryaa:mCherry for screening of transgenics. Clones were tested for correct recombination by digestion with restriction enzymes. The transgenic line was generated by injecting 10-20 pg of the final UAS:mem-Scarlet vector with 50 pg *tol2* transposase into nacre^−/−^ zebrafish eggs at the one cell stage. Founder animals were identified by outcrossing and screening for the red eye marker, then crossed with several Gal4 transgenic lines to confirm mem-Scarlet expression to select the best founder. All experiments were performed with the F2 and F3 generations. Multiple insertions were present in these generations.

#### Live-imaging

Larvae were anaesthetised with 600 µM tricaine in E3 embryo medium and immobilised in 1.5% low melting-point agarose on a glass coverslip, which was suspended over a microscope slide using high vacuum silicone grease to create a well containing E3 embryo medium and 600 µM tricaine. Z-stacks (with optimal z-step) were obtained using a Zeiss LSM880 microscope with Airyscan FAST in super-resolution mode, using a 20X objective lens (Zeiss Plan-Apochromat 20x dry, NA = 0.8). All Mauthner, MiMi, and Mid3i axon images were taken from a lateral view of the spinal cord centred around somite 15. Mauthner, MiMi, and Mid3i cell bodies were imaged from the dorsal surface of the hindbrain. All lateral view images depict anterior on the left and dorsal of the top, while dorsal view images depict anterior on the top. Figure panels were prepared using Fiji and Adobe Illustrator CS6.

#### Quantification of Axon Diameter

Axon diameter from AiryScan FAST confocal images were measured using the following scripts, which were written in ImageJ Macro Language using the open source image analysis software Fiji ^76, 77^.

A “Split Axons Tool” was produced to allow the splitting of two adjacent Mauthner (or MiMi/Mid3i axons) in whole larvae z-stack datasets. To achieve this, the macro rotates the dataset to give the top down (x-z plane) view, performs a maximum intensity projection and presents this to the user and asks for a point to be placed between the two axons. The macro then samples the image repeatedly along the z-axis in user configurable x-widths, to identify local minima between peak fluorescence. By doing so, the fluorescence trough between axons is mapped. The minima closest to the user selected point is selected and used to create two separate selections on duplicates of the original x-z rotated z-stack, for the upper and lower Mauthner axons, where the pixels outside the selections for the respective axons is set to zero grey values, before rotating these new z-stacks back to the front facing x-y plane for saving and analysis.

In order to measure axon diameter, two separate ImageJ tools were generated. Firstly, the “Axon Trace Tool” was written to trace the approximate centre point of the axon along its length. To do this, the edges of the axon are identified by forming an intensity profile along y-axis profile using the Plot Profile function in ImageJ (with a user configurable thickness in the x-axis) and the centre point for this location on the x-axis, given by the midpoint of the axon edges. This is then repeated along the x-axis to give a trace of the centre point of the axon for any given x coordinate.

Secondly, the “Axon Caliber Tool” was written to measure the average axon diameter along the length of the selected axon, and was based on the method used to measure axon diameter in ^7^. This macro takes the axon trace produced by the Axon Trace Tool, and for each location along the x-axis calculates the trajectory of the axon, based on a user configurable number of x-y centre coordinates before and after the current location. The macro then plots a line (of user configurable width) perpendicular to calculated trajectory and extracts the fluorescence profile along this line. The edges of the axon are then extracted from this profile, giving the width at this location along the x-axis. The macro then repeats this process along the x-axis and provides the diameter at each x-y coordinate, as well as the average thickness of the axon. Unless otherwise indicated, axon diameter was measured for only one axon per fish, selecting the axon located closest to the imaging objective.

#### Quantification of cell body size

Imaging was performed on *ipo13b^ue^*^57^; Tg(hspGFF62A:Gal4); Tg(UAS:mem-Scarlet) fish. The Mauthner neuron was imaged from the dorsal surface of the zebrafish hindbrain at 4 dpf. To delineate a boundary between the lateral dendrite and cell body, an ellipse was fit to the Mauthner cell body. The area of the ellipse was used as the measurement of cell body size.

#### Transmission electron microscopy

Zebrafish tissue was prepared for transmission electron microscopy according to a previously published protocol^78^. Briefly, zebrafish embryos were terminally anaesthetised in tricaine, heads removed for genotyping, and trunks incubated with microwave stimulation in a primary fixative of 4% paraformaldehyde + 2% glutaraldehyde in 0.1 M sodium cacodylate buffer. Tissue was then stored up to 3 weeks in primary fixative at 4°C. Samples were washed in 0.1 M Cacodylate buffer and incubated with microwave stimulation in secondary fixative of 2% osmium tetroxide in 0.1 M sodium cacodylate/0.1 M imidazole buffer pH7.5, then left overnight at room temperature. Samples were washed with distilled water, then stained en bloc with a saturated (8%) uranyl acetate solution with microwave stimulation followed by 1.5 hr incubation at room temperature. Next, samples were dehydrated with an ethanol series and acetone using microwave stimulation. Samples were embedded in Embed-812 resin (Electron Microscopy Sciences) and silver sections (70-80 nm in thickness) cut using a Reichert Jung Ultracut Microtome (Leica) and a Diatome diamond knife. Sections were mounted hexagonal copper electron microscopy grids (200 Mesh Grids, Agar Scientific). Mounted sections were stained in uranyl acetate and Sato lead stain. TEM imaging was performed at the University of Edinburgh Biology Scanning Electron Microscope Facility using a Jeol JEM1400 Plus Transmission Electron Microscope. Dorsal and ventral spinal cord were imaged at 12000x magnification and Mauthner axon at 15000x magnification. In order to create panoramic views, individual electron micrograph tiles were aligned using the automated photomerge tool in Adobe Photoshop CC. To assess axon diameter, axonal areas were measured by tracing axons in Fiji, which were then used to calculate diameter using the equation diameter = 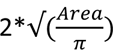.

#### Neurofilament Analysis

ROIs of 430 pixels x 430 pixels which were absent of mitochondria or other organelles were selected from each electron micrograph of the Mauthner axon for quantification. Neurofilaments in electron micrographs were detected using the ML-segmenter algorithm in Arivis Vision4D. To train the model in the algorithm, neurofilament and non-neurofilament class examples were annotated using a brush tool across 5 images. The trained model was used to segment neurofilament objects across the entire ROI image batch. Coordinates of neurofilament objects were exported and used in a custom Wolfram Mathematica 13.0 notebook to calculate the nearest neighbours using the built-in Nearest[] function. Neurofilament density was calculated using the number of neurofilaments within the ROI.

#### Cell-type specific targeting of ipo13 in neurons

Cas9 and sgRNA expression vectors were generated using a previously described strategy and protocol ^79, 80^. Firstly, a U6-based sgRNA construct expressing 3 sgRNAs targeting *ipo13b* was generated using Golden gate cloning. The following sgRNA sequences were used, which were designed using the CRISPRscan tool ^81^ and selected based on low prediction for off-targeting : *ipo13b* sgRNA1: GGTGCCTGAGGCCTGGCCGG (exon 3), *ipo13b* sgRNA2: GGTGCTGACGGTTCTTCCAG (exon 3), *ipo13b* sgRNA3: GGTTGAGTCGAGGACTACAG (exon 21). Each sgRNA was tested for efficiency prior to cloning by using them to generate F0 generation mutants (crispants), which phenocopied the *ipo13b^ue^*^57^ mutant phenotype. To clone the sgRNA sequences into the U6 expression vectors, annealed oligos containing the sgRNA and BsmBI sequences were generated using the following primer pairs: TTCGGTGCCTGAGGCCTGGCCGG & AAACCCGGCCAGGCCTCAGGCAC, TTCGGTGCTGACGGTTCTTCCAG & AAACCTGGAAGAACCGTCAGCAC, TTCGGGAGTCAAAGACATTGTGA & AAACTCACAATGTCTTTGACTCC. 200 µM of each oligo were mixed together in a 20 µl reaction with NEB Buffer 2.1 and incubated at 95°C for 5 min, then decreased to 50°C at 0.1°C/s, followed by a 10 min incubation at 50°C. The annealed oligos were then ligated into U6 promotor-based cassettes: *ipo13b* sgRNA1 into pU6a:sgRNA#1 (Addgene #64245), *ipo13b* sgRNA2 into pU6a:sgRNA#2 (Addgene #64246), and *ipo13b* sgRNA3 into pU6b:sgRNA#3 (Addgene #64247). Ligation reactions were carried out by mixing 1 µL 10x NEB CutSmart buffer, 1 µL T4 DNA ligase buffer, 100 ng U6 plasmid, 1 µl annealed oligos, 0.3 µL T4 DNA ligase, 0.3 µL BsmBI, 0.2 µl PstI, and 0.2 µL SalI in a 10 µL reaction, then incubating for 3 cycles of 37°C for 20 min and 16°C for 15 min. This was followed by 37°C for 10 min, 55°C for 15 min and 80°C for 15 min.

To construct the final sgRNA expressing vector containing all 3 sgRNAs, 50ng of pGGDestTol2LC-3sgRNA (Addgene #64241) was combined with 100 ng of each pU6:sgRNA vector generated above, 2 µL 10x NEB CutSmart Buffer, 2 µL T4 DNA ligase buffer, 1 µL T4 DNA ligase, and 1 µL BsaI in a 20 µL reaction. This was incubated with 3 cycles of 37°C for 20 min and 16°C for 15 min, followed by 15 min at 80°C. The destination vector of pGGDestTol2LC-3sgRNA contains a cerulean eye marker (cryaa:cerulean) for screening of transgenics. The sequence of the final expression vector was confirmed by sequencing before use.

To generate the construct expressing cas9 in neurons the Tol2-based multisite Gateway system was used. 20 fmol of each of the following entry and destination vectors were combined in a LR reaction using LR Clonase II Plus performed overnight at room temperature: p5E_nbt, pME_cas9 (Addgene #64237), p3E-polyA (Tol2Kit #302), and pDestTol2pA2-cryaa:mCherry (Addgene #64023). Note, the destination vector includes the red eye marker cryaa:mCherry for screening of transgenics. Clones were tested for correct recombination by digestion with restriction enzymes.

Two transgenic lines, Tg(U6:3sgRNA-ipo13b) and Tg(nbt:cas9), were generated by injecting 10-20 pg of the expression vector with 50 pg *tol2* transposase into Tg(mbp:eGFPCAAX) zebrafish eggs at the one cell stage. Founder animals were identified by outcrossing and screening for the red or blue eye markers. Founders were outcrossed to Tg(hspGFF62A:Gal4); Tg(UAS:RFP) to generate F1 offspring that carried either Tg(U6:3sgRNA-IPO13) or Tg(nbt:cas9), as well as Tg(hspGFF62A:Gal4); Tg(UAS:RFP) and Tg(mbp:eGFPCAAX) to label the Mauthner axon and myelin. Neuronal specific mutants were generated by crossing Tg(U6:3sgRNA-ipo13b) and Tg(nbt:cas9) lines and screening for the presence of both red and cerulean eye markers. All experiments were performed using parents from the F1 and F2 generations. Multiple insertions were present in these generations.

#### Electrophysiology

Zebrafish were dissected as described previously ^82^ to access the Mauthner neuron. For experiments using neuronal specific *ipo13b* mutants, fish were pre-screened and only those exhibiting reduced axon diameter were used for electrophysiology. In short, 2-5 dpf anaesthetised zebrafish were laid on their sides on a Sylgard dish and secured using tungsten pins through their notochords in a dissection solution containing the following: 134 mM NaCl, 2.9 KCl mM, 2.1 CaCl_2_ mM, 1.2 MgCl_2_ mM, 10 mM HEPES, 10 mM Glucose and 600 µM tricaine, adjusted to pH 7.8 with NaOH. Their eyes and lower and upper jaws were removed using forceps to expose the ventral surface of the hindbrain, which was secured with an additional tungsten pin. The Mauthner axons was exposed, alongside motor neurons, as described by ^83^. A dissecting tungsten pin was used to remove the skin and the muscle overlaying the Mauthner axons and motoneurons in a single segment. Following the dissection, zebrafish together with their recording chamber were moved to the recording rig and washed with extracellular solution containing the following: 134 mM NaCl, 2.9 mM KCl, 2.1 mM CaCl_2_, 1.2 mM MgCl_2_, 10 mM HEPES, 10 mM Glucose and 15 µM Tubocurarine. The cells were visualised using an Olympus microscope capable of Differential Interference Constrast (DIC) using a 60X water immersion, NA=1 objective lens and a ROLERA Bolt sCMOS camera (QImaging) with Q capture software. The stimulating electrode filled with extracellular solution was then positioned in the spinal cord touching the exposed neurons underneath. Mauthner whole-cell recordings were performed with thick-walled borosilicate glass pipettes pulled to 6–10 MΩ. The internal solution contained the following: 125 mM K-gluconate, 15 mM KCl, 10 mM HEPES, 5 mM EGTA, 2 mM MgCl_2_, 0.4 mM NaGTP, 2 mM NaATP, and 10 mM Na-phosphocreatine, adjusted to pH 7.4 with KOH. A 270s recording was performed in a current clamp configuration, and the cell resting membrane potential was established as an average of the first 10 seconds of the recording if the cell did not fire during that time. To measure the conduction velocity along the Mauthner axon, 270s after breaking through the cell the zebrafish were washed with recording solution containing the following: 134 mM NaCl, 2.9 mM KCl, 2.1 mM CaCl_2_, 1.2 mM MgCl_2_, 10 mM HEPES, 10 mM Glucose. Blockers of fast synaptic transmission were supplemented to this solution: 50 µM AP5, 20 µM strychnine, 100 µM picrotoxin and 50 µM CNQX. The antidromic Mauthner action potentials were recorded following the field stimulation by the stimulating electrode connected to DS2A Isolated Voltage Stimulator (Digitimer) in the spinal cord. The action potentials were recoded using Clampex 10.6 (Molecular Devices) at 100 kHz sampling rate and filtered at 2 kHz using a MultiClamp 700B amplifier (Molecular Devices). 30 consecutive action potentials were recoded every 5 seconds using Clampex 10.6 (Molecular Devices) at 100 kHz sampling rate and filtered at 2 kHz using a MultiClamp 700B amplifier (Molecular Devices). At the end of the recording, images of the zebrafish were obtained with a 4X objective and stitched using Adobe Photoshop. The resulting image was then transferred to FIJI and the distance between the stimulating and recording electrode was measured. The latency of the action potential peak was calculated by the in-house built Matlab Script ^52^ available at https://github.com/skotuke/Mauthner_analysis. The conduction velocity of action potential was calculated dividing the distance between the electrodes by the latency from action potential peak to the stimulus artifact. For the analysis of action potential fidelity consecutive trains of 10 stimuli were delivered at 1, 10, 100, 300, 500 and 1000Hz every 30s. Recordings were made at 20 kHz sampling rate and filtered at 2 kHz. The number of action potentials were calculated using Clampfit software and the action potential success rate was calculated as a number of action potentials fired out of 10. The precision of action potential arrival was calculated as the standard deviation of 30 consecutive action potential latencies from the stimulus artifact.

#### Behavioural Analysis

At 4 dpf individual larvae were added to each well of a 24 well plate in 1.5 mL of 10 mM HEPES-buffered E3 embryo medium. The next day, a 20 min period of free swimming was recorded using a DanioVision Box and Ethovision XT 14 software (Noldus Information Technology, Netherlands). Ethovision XT 14 software was used to determine the percentage of time spent moving. A track smoothing profile was added setting the minimal distance moved to > 0.20 mm and the maximum distance moved to > 9.20 mm. Movement was calculated using an averaging interval of 3 samples, start velocity of 5.00 mm/s and stop velocity of 1.00 mm/s. Only fish with inflated swim bladders were analyzed.

### QUANTIFICATION AND STATISTICAL ANALYSIS

Statistical analysis was performed using GraphPad Prism 8 software. Unless otherwise indicated, n number represents individual fish (or a single axon from an individual fish). All data are presented as mean +/− standard deviation, aside from behavioural data which is presented as box and whisker plots indicating the median, 25^th^ and 75^th^ percentiles, max and min. N values, p values and statistical tests are noted in the figure legends. Significance was defined as p<0.05.

## References

1. Aboitiz, F., Scheibel, A.B., Fisher, R.S., and Zaidel, E. (1992). Fiber composition of the human corpus callosum. Brain Res 598, 143–153. 10.1016/0006-8993(92)90178-c.

2. Terao, S., Sobue, G., Hashizume, Y., Shimada, N., and Mitsuma, T. (1994). Age-related changes of the myelinated fibers in the human corticospinal tract: a quantitative analysis. Acta Neuropathol 88, 137–142. 10.1007/BF00294506.

3. Perge, J.A., Niven, J.E., Mugnaini, E., Balasubramanian, V., and Sterling, P. (2012). Why do axons differ in caliber? J Neurosci 32, 626–638. 10.1523/JNEUROSCI.4254-11.2012.

4. Seidl, A.H., and Rubel, E.W. (2016). Systematic and differential myelination of axon collaterals in the mammalian auditory brainstem. Glia 64, 487–494. 10.1002/glia.22941.

5. Pesaresi, M., Soon-Shiong, R., French, L., Kaplan, D.R., Miller, F.D., and Paus, T. (2015). Axon diameter and axonal transport: In vivo and in vitro effects of androgens. Neuroimage 115, 191–201. 10.1016/j.neuroimage.2015.04.048.

6. Genc, S., Raven, E.P., Drakesmith, M., Blakemore, S.J., and Jones, D.K. (2023). Novel insights into axon diameter and myelin content in late childhood and adolescence. Cereb Cortex. 10.1093/cercor/bhac515.

7. Chereau, R., Saraceno, G.E., Angibaud, J., Cattaert, D., and Nagerl, U.V. (2017). Superresolution imaging reveals activity-dependent plasticity of axon morphology linked to changes in action potential conduction velocity. Proc Natl Acad Sci U S A 114, 1401–1406. 10.1073/pnas.1607541114.

8. Sinclair, J.L., Fischl, M.J., Alexandrova, O., Hebeta, M., Grothe, B., Leibold, C., and Kopp-Scheinpflug, C. (2017). Sound-Evoked Activity Influences Myelination of Brainstem Axons in the Trapezoid Body. J Neurosci 37, 8239–8255. 10.1523/JNEUROSCI.3728-16.2017.

9. Nicholson, M., Wood, R.J., Gonsalvez, D.G., Hannan, A.J., Fletcher, J.L., Xiao, J., and Murray, S.S. (2022). Remodelling of myelinated axons and oligodendrocyte differentiation is stimulated by environmental enrichment in the young adult brain. Eur J Neurosci 56, 6099–6114. 10.1111/ejn.15840.

10. Bechler, M.E., Byrne, L., and Ffrench-Constant, C. (2015). CNS Myelin Sheath Lengths Are an Intrinsic Property of Oligodendrocytes. Curr Biol 25, 2411–2416. 10.1016/j.cub.2015.07.056.

11. Goebbels, S., Wieser, G.L., Pieper, A., Spitzer, S., Weege, B., Yan, K., Edgar, J.M., Yagensky, O., Wichert, S.P., Agarwal, A., et al. (2017). A neuronal PI(3,4,5)P3-dependent program of oligodendrocyte precursor recruitment and myelination. Nat Neurosci 20, 10-15. 10.1038/nn.4425.

12. Lee, S., Leach, M.K., Redmond, S.A., Chong, S.Y., Mellon, S.H., Tuck, S.J., Feng, Z.Q., Corey, J.M., and Chan, J.R. (2012). A culture system to study oligodendrocyte myelination processes using engineered nanofibers. Nat Methods 9, 917–922. 10.1038/nmeth.2105.

13. Mayoral, S.R., Etxeberria, A., Shen, Y.A., and Chan, J.R. (2018). Initiation of CNS Myelination in the Optic Nerve Is Dependent on Axon Caliber. Cell Rep 25, 544–550 e543. 10.1016/j.celrep.2018.09.052.

14. Stedehouder, J., Brizee, D., Slotman, J.A., Pascual-Garcia, M., Leyrer, M.L., Bouwen, B.L., Dirven, C.M., Gao, Z., Berson, D.M., Houtsmuller, A.B., and Kushner, S.A. (2019). Local axonal morphology guides the topography of interneuron myelination in mouse and human neocortex. Elife 8. 10.7554/eLife.48615.

15. Voyvodic, J.T. (1989). Target size regulates calibre and myelination of sympathetic axons. Nature 342, 430–433. 10.1038/342430a0.

16. Waxman, S.G. (1980). Determinants of conduction velocity in myelinated nerve fibers. Muscle Nerve 3, 141–150. 10.1002/mus.880030207.

17. Golden, C.E.M., Yee, Y., Wang, V.X., Harony-Nicolas, H., Hof, P.R., Lerch, J.P., and Buxbaum, J.D. (2020). Reduced axonal caliber and structural changes in a rat model of Fragile X syndrome with a deletion of a K-Homology domain of Fmr1. Transl Psychiatry 10, 280. 10.1038/s41398-020-00943-x.

18. Huang, S.Y., Fan, Q., Machado, N., Eloyan, A., Bireley, J.D., Russo, A.W., Tobyne, S.M., Patel, K.R., Brewer, K., Rapaport, S.F., et al. (2019). Corpus callosum axon diameter relates to cognitive impairment in multiple sclerosis. Ann Clin Transl Neurol 6, 882–892. 10.1002/acn3.760.

19. Klok, M.D., Bugiani, M., de Vries, S.I., Gerritsen, W., Breur, M., van der Sluis, S., Heine, V.M., Kole, M.H.P., Baron, W., and van der Knaap, M.S. (2018). Axonal abnormalities in vanishing white matter. Ann Clin Transl Neurol 5, 429–444. 10.1002/acn3.540.

20. Liu, M., Xu, P., Guan, Z., Qian, X., Dockery, P., Fitzgerald, U., O’Brien, T., and Shen, S. (2018). Ulk4 deficiency leads to hypomyelination in mice. Glia 66, 175–190. 10.1002/glia.23236.

21. Lousada, E., Boudreau, M., Cohen-Adad, J., Nait Oumesmar, B., Burguiere, E., and Schreiweis, C. (2021). Reduced Axon Calibre in the Associative Striatum of the Sapap3 Knockout Mouse. Brain Sci 11. 10.3390/brainsci11101353.

22. Richards, K., Jancovski, N., Hanssen, E., Connelly, A., and Petrou, S. (2021). Atypical myelinogenesis and reduced axon caliber in the Scn1a variant model of Dravet syndrome: An electron microscopy pilot study of the developing and mature mouse corpus callosum. Brain Res 1751, 147157. 10.1016/j.brainres.2020.147157.

23. Wegiel, J., Kaczmarski, W., Flory, M., Martinez-Cerdeno, V., Wisniewski, T., Nowicki, K., Kuchna, I., and Wegiel, J. (2018). Deficit of corpus callosum axons, reduced axon diameter and decreased area are markers of abnormal development of interhemispheric connections in autistic subjects. Acta Neuropathol Commun 6, 143. 10.1186/s40478-018-0645-7.

24. Xie, M., Tobin, J.E., Budde, M.D., Chen, C.I., Trinkaus, K., Cross, A.H., McDaniel, D.P., Song, S.K., and Armstrong, R.C. (2010). Rostrocaudal analysis of corpus callosum demyelination and axon damage across disease stages refines diffusion tensor imaging correlations with pathological features. J Neuropathol Exp Neurol 69, 704–716. 10.1097/NEN.0b013e3181e3de90.

25. Fan, A., Tofangchi, A., Kandel, M., Popescu, G., and Saif, T. (2017). Coupled circumferential and axial tension driven by actin and myosin influences in vivo axon diameter. Sci Rep 7, 14188. 10.1038/s41598-017-13830-1.

26. Friede, R.L., and Samorajski, T. (1970). Axon caliber related to neurofilaments and microtubules in sciatic nerve fibers of rats and mice. Anat Rec 167, 379–387. 10.1002/ar.1091670402.

27. Leite, S.C., Sampaio, P., Sousa, V.F., Nogueira-Rodrigues, J., Pinto-Costa, R., Peters, L.L., Brites, P., and Sousa, M.M. (2016). The Actin-Binding Protein alpha-Adducin Is Required for Maintaining Axon Diameter. Cell Rep 15, 490–498. 10.1016/j.celrep.2016.03.047.

28. Ohara, O., Gahara, Y., Miyake, T., Teraoka, H., and Kitamura, T. (1993). Neurofilament deficiency in quail caused by nonsense mutation in neurofilament-L gene. J Cell Biol 121, 387–395. 10.1083/jcb.121.2.387.

29. Stephan, R., Goellner, B., Moreno, E., Frank, C.A., Hugenschmidt, T., Genoud, C., Aberle, H., and Pielage, J. (2015). Hierarchical microtubule organization controls axon caliber and transport and determines synaptic structure and stability. Dev Cell 33, 5–21. 10.1016/j.devcel.2015.02.003.

30. Garcia, M.L., Rao, M.V., Fujimoto, J., Garcia, V.B., Shah, S.B., Crum, J., Gotow, T., Uchiyama, Y., Ellisman, M., Calcutt, N.A., and Cleveland, D.W. (2009). Phosphorylation of highly conserved neurofilament medium KSP repeats is not required for myelin-dependent radial axonal growth. J Neurosci 29, 1277–1284. 10.1523/JNEUROSCI.3765-08.2009.

31. Garcia, M.L., Lobsiger, C.S., Shah, S.B., Deerinck, T.J., Crum, J., Young, D., Ward, C.M., Crawford, T.O., Gotow, T., Uchiyama, Y., et al. (2003). NF-M is an essential target for the myelin-directed "outside-in" signaling cascade that mediates radial axonal growth. J Cell Biol 163, 1011–1020. 10.1083/jcb.200308159.

32. Suminaite, D., Lyons, D.A., and Livesey, M.R. (2019). Myelinated axon physiology and regulation of neural circuit function. Glia 67, 2050–2062. 10.1002/glia.23665.

33. Quan, Y., Ji, Z.L., Wang, X., Tartakoff, A.M., and Tao, T. (2008). Evolutionary and transcriptional analysis of karyopherin beta superfamily proteins. Mol Cell Proteomics 7, 1254–1269. 10.1074/mcp.M700511-MCP200.

34. Kegel, L., Rubio, M., Almeida, R.G., Benito, S., Klingseisen, A., and Lyons, D.A. (2019). Forward Genetic Screen Using Zebrafish to Identify New Genes Involved in Myelination. Methods Mol Biol 1936, 185–209. 10.1007/978-1-4939-9072-6_11.

35. Klingseisen, A., Ristoiu, A.M., Kegel, L., Sherman, D.L., Rubio-Brotons, M., Almeida, R.G., Koudelka, S., Benito-Kwiecinski, S.K., Poole, R.J., Brophy, P.J., and Lyons, D.A. (2019). Oligodendrocyte Neurofascin Independently Regulates Both Myelin Targeting and Sheath Growth in the CNS. Dev Cell 51, 730–744 e736. 10.1016/j.devcel.2019.10.016.

36. Marshall-Phelps, K.L.H., Kegel, L., Baraban, M., Ruhwedel, T., Almeida, R.G., Rubio-Brotons, M., Klingseisen, A., Benito-Kwiecinski, S.K., Early, J.J., Bin, J.M., et al. (2020). Neuronal activity disrupts myelinated axon integrity in the absence of NKCC1b. J Cell Biol 219. 10.1083/jcb.201909022.

37. Mingot, J.M., Kostka, S., Kraft, R., Hartmann, E., and Gorlich, D. (2001). Importin 13: a novel mediator of nuclear import and export. EMBO J 20, 3685–3694. 10.1093/emboj/20.14.3685.

38. Kimura, M., and Imamoto, N. (2014). Biological significance of the importin-beta family-dependent nucleocytoplasmic transport pathways. Traffic 15, 727–748. 10.1111/tra.12174.

39. Sachse, S.M., Lievens, S., Ribeiro, L.F., Dascenco, D., Masschaele, D., Horre, K., Misbaer, A., Vanderroost, N., De Smet, A.S., Salta, E., et al. (2019). Nuclear import of the DSCAM-cytoplasmic domain drives signaling capable of inhibiting synapse formation. EMBO J 38. 10.15252/embj.201899669.

40. Panayotis, N., Karpova, A., Kreutz, M.R., and Fainzilber, M. (2015). Macromolecular transport in synapse to nucleus communication. Trends Neurosci 38, 108–116. 10.1016/j.tins.2014.12.001.

41. Lever, M.B., Karpova, A., and Kreutz, M.R. (2015). An Importin Code in neuronal transport from synapse-to-nucleus? Front Mol Neurosci 8, 33. 10.3389/fnmol.2015.00033.

42. Rishal, I., and Fainzilber, M. (2014). Axon-soma communication in neuronal injury. Nat Rev Neurosci 15, 32–42. 10.1038/nrn3609.

43. Bono, F., Cook, A.G., Grunwald, M., Ebert, J., and Conti, E. (2010). Nuclear import mechanism of the EJC component Mago-Y14 revealed by structural studies of importin 13. Mol Cell 37, 211–222. 10.1016/j.molcel.2010.01.007.

44. Perrot, R., Berges, R., Bocquet, A., and Eyer, J. (2008). Review of the multiple aspects of neurofilament functions, and their possible contribution to neurodegeneration. Mol Neurobiol 38, 27–65. 10.1007/s12035-008-8033-0.

45. Zhang, Y., Chen, K., Sloan, S.A., Bennett, M.L., Scholze, A.R., O’Keeffe, S., Phatnani, H.P., Guarnieri, P., Caneda, C., Ruderisch, N., et al. (2014). An RNA-sequencing transcriptome and splicing database of glia, neurons, and vascular cells of the cerebral cortex. J Neurosci 34, 11929–11947. 10.1523/JNEUROSCI.1860-14.2014.

46. Brady, S.T., Witt, A.S., Kirkpatrick, L.L., de Waegh, S.M., Readhead, C., Tu, P.H., and Lee, V.M. (1999). Formation of compact myelin is required for maturation of the axonal cytoskeleton. J Neurosci 19, 7278–7288.

47. de Waegh, S.M., Lee, V.M., and Brady, S.T. (1992). Local modulation of neurofilament phosphorylation, axonal caliber, and slow axonal transport by myelinating Schwann cells. Cell 68, 451–463. 10.1016/0092-8674(92)90183-d.

48. Eichel, M.A., Gargareta, V.I., D’Este, E., Fledrich, R., Kungl, T., Buscham, T.J., Luders, K.A., Miracle, C., Jung, R.B., Distler, U., et al. (2020). CMTM6 expressed on the adaxonal Schwann cell surface restricts axonal diameters in peripheral nerves. Nat Commun 11, 4514. 10.1038/s41467-020-18172-7.

49. Sanchez, I., Hassinger, L., Paskevich, P.A., Shine, H.D., and Nixon, R.A. (1996). Oligodendroglia regulate the regional expansion of axon caliber and local accumulation of neurofilaments during development independently of myelin formation. J Neurosci 16, 5095–5105.

50. Yin, X., Crawford, T.O., Griffin, J.W., Tu, P., Lee, V.M., Li, C., Roder, J., and Trapp, B.D. (1998). Myelin-associated glycoprotein is a myelin signal that modulates the caliber of myelinated axons. J Neurosci 18, 1953–1962.

51. Hutchinson, N.A., Koles, Z.J., and Smith, R.S. (1970). Conduction velocity in myelinated nerve fibres of Xenopus laevis. J Physiol 208, 279–289. 10.1113/jphysiol.1970.sp009119.

52. Madden, M.E., Suminaite, D., Ortiz, E., Early, J.J., Koudelka, S., Livesey, M.R., Bianco, I.H., Granato, M., and Lyons, D.A. (2021). CNS Hypomyelination Disrupts Axonal Conduction and Behavior in Larval Zebrafish. J Neurosci 41, 9099–9111. 10.1523/JNEUROSCI.0842-21.2021.

53. Gallent, E.A., and Steward, O. (2018). Neuronal PTEN deletion in adult cortical neurons triggers progressive growth of cell bodies, dendrites, and axons. Exp Neurol 303, 12–28. 10.1016/j.expneurol.2018.01.005.

54. LaSarge, C.L., Santos, V.R., and Danzer, S.C. (2015). PTEN deletion from adult-generated dentate granule cells disrupts granule cell mossy fiber axon structure. Neurobiol Dis 75, 142–150. 10.1016/j.nbd.2014.12.029.

55. Markus, A., Zhong, J., and Snider, W.D. (2002). Raf and akt mediate distinct aspects of sensory axon growth. Neuron 35, 65–76. 10.1016/s0896-6273(02)00752-3.

56. Baade, I., Spillner, C., Schmitt, K., Valerius, O., and Kehlenbach, R.H. (2018). Extensive Identification and In-depth Validation of Importin 13 Cargoes. Mol Cell Proteomics 17, 1337–1353. 10.1074/mcp.RA118.000623.

57. Fatima, S., Wagstaff, K.M., Lieu, K.G., Davies, R.G., Tanaka, S.S., Yamaguchi, Y.L., Loveland, K.L., Tam, P.P., and Jans, D.A. (2016). Interactome of the inhibitory isoform of the nuclear transporter Importin 13. Biochim Biophys Acta 1864, 546–561. 10.1016/j.bbamcr.2016.12.017.

58. Kimura, M., Morinaka, Y., Imai, K., Kose, S., Horton, P., and Imamoto, N. (2017). Extensive cargo identification reveals distinct biological roles of the 12 importin pathways. Elife 6. 10.7554/eLife.21184.

59. Mackmull, M.T., Klaus, B., Heinze, I., Chokkalingam, M., Beyer, A., Russell, R.B., Ori, A., and Beck, M. (2017). Landscape of nuclear transport receptor cargo specificity. Mol Syst Biol 13, 962. 10.15252/msb.20177608.

60. Giagtzoglou, N., Lin, Y.Q., Haueter, C., and Bellen, H.J. (2009). Importin 13 regulates neurotransmitter release at the Drosophila neuromuscular junction. J Neurosci 29, 5628–5639. 10.1523/JNEUROSCI.0794-09.2009.

61. Xin, W., and Chan, J.R. (2020). Myelin plasticity: sculpting circuits in learning and memory. Nat Rev Neurosci 21, 682–694. 10.1038/s41583-020-00379-8.

62. Almeida, R.G., and Lyons, D.A. (2017). On Myelinated Axon Plasticity and Neuronal Circuit Formation and Function. J Neurosci 37, 10023–10034. 10.1523/JNEUROSCI.3185-16.2017.

63. Bonetto, G., Kamen, Y., Evans, K.A., and Karadottir, R.T. (2020). Unraveling Myelin Plasticity. Front Cell Neurosci 14, 156. 10.3389/fncel.2020.00156.

64. Lister, J.A., Robertson, C.P., Lepage, T., Johnson, S.L., and Raible, D.W. (1999). nacre encodes a zebrafish microphthalmia-related protein that regulates neural-crest-derived pigment cell fate. Development 126, 3757–3767.

65. Almeida, R.G., Czopka, T., Ffrench-Constant, C., and Lyons, D.A. (2011). Individual axons regulate the myelinating potential of single oligodendrocytes in vivo. Development 138, 4443–4450. 10.1242/dev.071001.

66. Asakawa, K., Suster, M.L., Mizusawa, K., Nagayoshi, S., Kotani, T., Urasaki, A., Kishimoto, Y., Hibi, M., and Kawakami, K. (2008). Genetic dissection of neural circuits by Tol2 transposon-mediated Gal4 gene and enhancer trapping in zebrafish. Proc Natl Acad Sci U S A 105, 1255–1260. 10.1073/pnas.0704963105.

67. Yamanaka, I., Miki, M., Asakawa, K., Kawakami, K., Oda, Y., and Hirata, H. (2013). Glycinergic transmission and postsynaptic activation of CaMKII are required for glycine receptor clustering in vivo. Genes Cells 18, 211–224. 10.1111/gtc.12032.

68. Kelsh, R.N., Brand, M., Jiang, Y.J., Heisenberg, C.P., Lin, S., Haffter, P., Odenthal, J., Mullins, M.C., van Eeden, F.J., Furutani-Seiki, M., et al. (1996). Zebrafish pigmentation mutations and the processes of neural crest development. Development 123, 369–389.

69. Minevich, G., Park, D.S., Blankenberg, D., Poole, R.J., and Hobert, O. (2012). CloudMap: a cloud-based pipeline for analysis of mutant genome sequences. Genetics 192, 1249–1269. 10.1534/genetics.112.144204.

70. Butler, M.G., Iben, J.R., Marsden, K.C., Epstein, J.A., Granato, M., and Weinstein, B.M. (2015). SNPfisher: tools for probing genetic variation in laboratory-reared zebrafish. Development 142, 1542–1552. 10.1242/dev.118786.

71. LaFave, M.C., Varshney, G.K., Vemulapalli, M., Mullikin, J.C., and Burgess, S.M. (2014). A defined zebrafish line for high-throughput genetics and genomics: NHGRI-1. Genetics 198, 167–170. 10.1534/genetics.114.166769.

72. Obholzer, N., Swinburne, I.A., Schwab, E., Nechiporuk, A.V., Nicolson, T., and Megason, S.G. (2012). Rapid positional cloning of zebrafish mutations by linkage and homozygosity mapping using whole-genome sequencing. Development 139, 4280–4290. 10.1242/dev.083931.

73. Shah, A.N., Davey, C.F., Whitebirch, A.C., Miller, A.C., and Moens, C.B. (2015). Rapid reverse genetic screening using CRISPR in zebrafish. Nat Methods 12, 535–540. 10.1038/nmeth.3360.

74. Dehairs, J., Talebi, A., Cherifi, Y., and Swinnen, J.V. (2016). CRISP-ID: decoding CRISPR mediated indels by Sanger sequencing. Sci Rep 6, 28973. 10.1038/srep28973.

75. Kwan, K.M., Fujimoto, E., Grabher, C., Mangum, B.D., Hardy, M.E., Campbell, D.S., Parant, J.M., Yost, H.J., Kanki, J.P., and Chien, C.B. (2007). The Tol2kit: a multisite gateway-based construction kit for Tol2 transposon transgenesis constructs. Dev Dyn 236, 3088–3099. 10.1002/dvdy.21343.

76. Schindelin, J., Arganda-Carreras, I., Frise, E., Kaynig, V., Longair, M., Pietzsch, T., Preibisch, S., Rueden, C., Saalfeld, S., Schmid, B., et al. (2012). Fiji: an open-source platform for biological-image analysis. Nat Methods 9, 676-682. 10.1038/nmeth.2019.

77. Schneider, C.A., Rasband, W.S., and Eliceiri, K.W. (2012). NIH Image to ImageJ: 25 years of image analysis. Nat Methods 9, 671–675. 10.1038/nmeth.2089.

78. Czopka, T., Ffrench-Constant, C., and Lyons, D.A. (2013). Individual oligodendrocytes have only a few hours in which to generate new myelin sheaths in vivo. Dev Cell 25, 599–609. 10.1016/j.devcel.2013.05.013.

79. Yin, L., Maddison, L.A., Li, M., Kara, N., LaFave, M.C., Varshney, G.K., Burgess, S.M., Patton, J.G., and Chen, W. (2015). Multiplex Conditional Mutagenesis Using Transgenic Expression of Cas9 and sgRNAs. Genetics 200, 431–441. 10.1534/genetics.115.176917.

80. Yin, L., Maddison, L.A., and Chen, W. (2016). Multiplex conditional mutagenesis in zebrafish using the CRISPR/Cas system. Methods Cell Biol 135, 3–17. 10.1016/bs.mcb.2016.04.018.

81. Moreno-Mateos, M.A., Vejnar, C.E., Beaudoin, J.D., Fernandez, J.P., Mis, E.K., Khokha, M.K., and Giraldez, A.J. (2015). CRISPRscan: designing highly efficient sgRNAs for CRISPR-Cas9 targeting in vivo. Nat Methods 12, 982–988. 10.1038/nmeth.3543.

82. Roy, B., and Ali, D.W. (2013). Patch clamp recordings from embryonic zebrafish Mauthner cells. J Vis Exp. 10.3791/50551.

83. Wen, H., and Brehm, P. (2005). Paired motor neuron-muscle recordings in zebrafish test the receptor blockade model for shaping synaptic current. J Neurosci 25, 8104–8111. 10.1523/JNEUROSCI.2611-05.2005.

